# Cell cycle gene regulation dynamics revealed by RNA velocity and deep-learning

**DOI:** 10.1101/2021.03.17.435887

**Authors:** Andrea Riba, Attila Oravecz, Matej Durik, Sara Jiménez, Violaine Alunni, Marie Cerciat, Matthieu Jung, Céline Keime, William M. Keyes, Nacho Molina

## Abstract

The cell cycle is a fundamental process of life, however, a quantitative understanding of gene regulation dynamics in the context of the cell cycle is still far from complete. Single-cell RNA-sequencing (scRNA-seq) technology gives access to its dynamics without externally perturbing the cell. Here, we build a high-resolution map of the cell cycle transcriptome based on scRNA-seq and deep-learning. By generating scRNA-seq libraries with high depth, in mouse embryonic stem cells and human fibroblasts, we are able to observe cycling patterns in the unspliced-spliced RNA space for single genes. Since existing methods in scRNA-seq are not efficient to measure cycling gene dynamics, we propose a deep learning approach to fit these cycling patterns sorting single cells across the cell cycle. We characterize the cell cycle in asynchronous pluripotent and differentiated cells identifying major waves of transcription during the G1 phase and systematically study the G1-G0 transition where the cells exit the cycle. Our work presents to the scientific community a broader understanding of RNA velocity and cell cycle maps, that we applied to pluripotency and differentiation. Our approach will facilitate the study of the cell cycle in multiple cellular models and different biological contexts, such as cancer and development.

## Introduction

Cells divide by progressing through highly organized phases in which they grow, synthesize a copy of their genetic material, and, finally, undergo mitosis^1^. Alternatively, cells can stop cycling and reversibly transition into quiescence or irreversibly differentiate or become senescent^2^. These processes require tight dynamic regulation of gene expression and despite immense research during the past decades, a quantitative picture of the gene regulation dynamics across the cell cycle is still incomplete. With the advent of single-cell RNA sequencing (scRNA-seq), scientists can now analyze intrinsically asynchronous cell-populations enabling the simultaneous identification of cells at different cell cycle stages. scRNA-seq provides a high-resolution approach to study the cell cycle without external perturbations, such as synchronization by drugs or engineered fluorescent reporters^3, 4^. Many attempts to computationally assign cell cycle phases have been performed^5–8^, but they typically lack generalizability and fail in capturing the correct cell cycle dynamics. Thanks to the depth of the scRNA-seq datasets generated in this paper, cycling patterns in the unspliced-spliced RNA space for single genes (RNA velocity) can be observed clearly and exploited to naturally sort cells across the cell cycle^9^. Combining RNA velocity with deep learning, we designed DeepCycle (https://github.com/andreariba/DeepCycle), a tool that assigns a continuous high-resolution cell cycle trajectory to single cells. The approach applies to different systems and has self-consistency checks to establish whether the analysis worked properly. DeepCycle allows us to estimate the dynamics of gene activation and deactivation with minimal assumptions, resulting in fits of gene expression kinetics. The fits naturally generate gene expression series that can be analyzed to obtain detailed kinetic parameters.

The link between cell cycle regulation and cell identity is not fully understood^10, 11^. Nonetheless, the cell cycle phases are deeply affected by the degree of stemness, for example, pluripotent and neural stem cells have short G1 phases, while committed cells extend their G1 phases and present overall longer cell cycles^11–13^. Thanks to DeepCycle, we not only recapitulate these findings in mESCs and human fibroblasts, but also extend the analysis to a public dataset of ductal cell progenitors, and uncover the underlying regulatory mechanisms, showing the different genes and transcription factors active in the different cellular models across the cell cycle.

Finally, as most of the cells within multicellular organisms are not actively cycling, tight control over cell cycle entry and exit is critical, as seen for example in organism development, hematopoiesis, activation of adaptive immune responses, and wound healing. However, in diseases like cancers, cells do not consistently respond to such regulatory cues and signals. It is thus important to understand the processes that determine cell cycle entry, cell cycle progression, and exit to quiescence^14^. Here, we characterize the branching point where human fibroblasts exit from the cell cycle. New marker genes and transcription factors underlying the process have been highlighted, paving the way to the systematic characterization of the G1-G0 transition in other cellular models.

## Results

### Generation of deep-sequenced single-cell RNA-seq datasets

To robustly study the cell cycle, we reasoned that the dataset should be enriched for growing-cycling cells. The majority of public scRNA-seq datasets have been generated to study the overall population of cells in a given condition, and, typically, they contain heterogeneous cell types. Therefore, we cultured mouse embryonic stem cells (mESCs) in 2i+LIF medium to maintain the ground state of pluripotency by blocking differentiation^15–17^, and generated an scRNA-seq library containing more than five thousand mESCs (Figure 1A). We included in our scRNA-seq analysis ductal cell progenitors from endocrine development (henceforth referred to as *ductal cells*) (Figure 1B)^18^. These cells have been linked to a proliferative cell state by specific marker genes^19^. Finally, to compare the results to a different cell type from another organism we also sequenced 5367 human fibroblasts (Figure 1C). These fibroblasts separate into two subpopulations, only one of which expressing cell cycle genes (Supplementary Figure S1, 16 out of the top 25 genes belong to the DAVID Keywords Cell cycle, Benjamini=2e-15), therefore we first focused on the proliferative subpopulation (n=3086).

**Figure 1.**
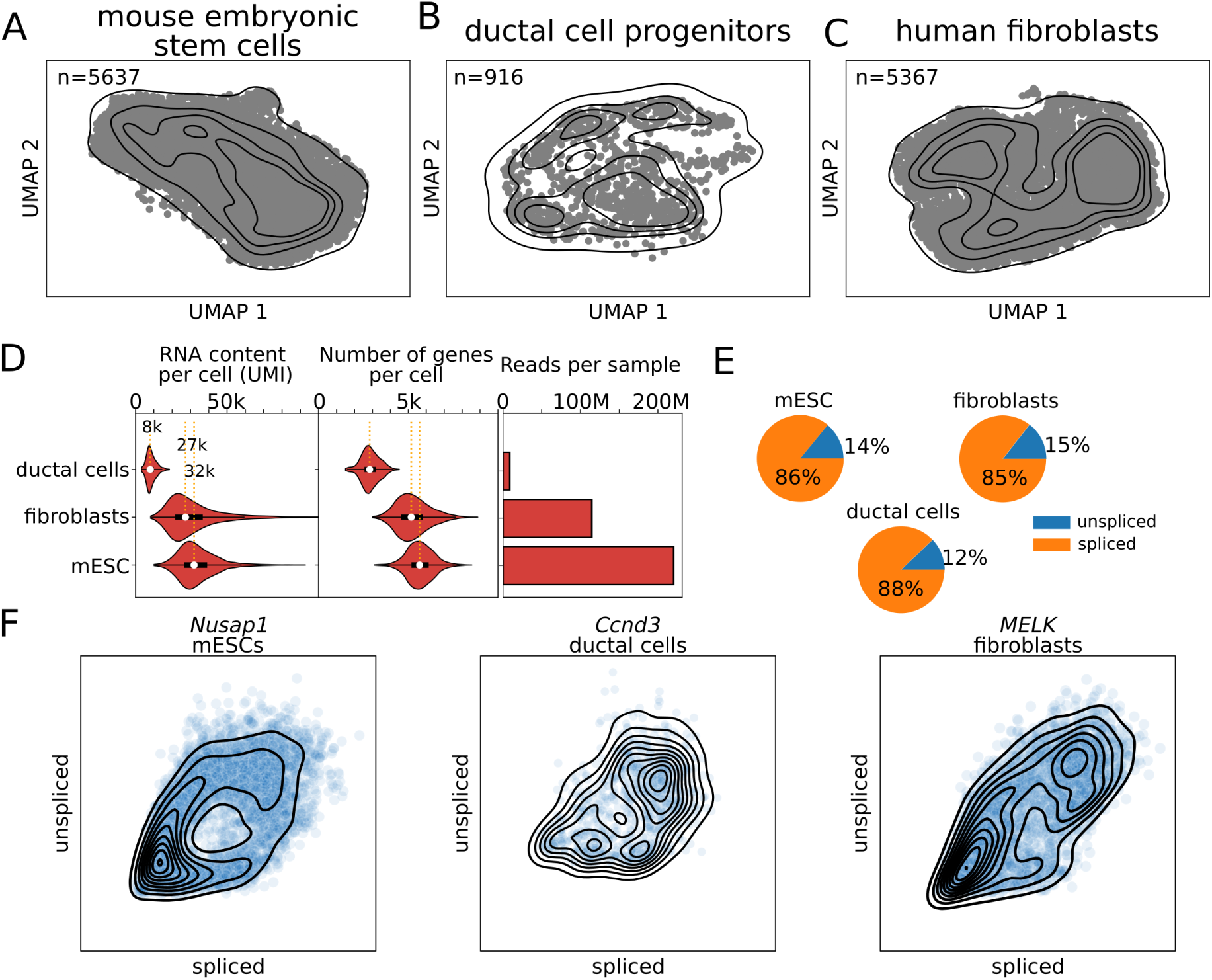
Single-cell RNA-sequencing data show the RNA velocity patterns. UMAP projections for mouse embryonic stem cells (**A**), ductal cell progenitors (**B**), and human fibroblasts (**C**). **D.** Distribution of RNA content (UMI), number of genes per cell, and the total number of reads for each sample. **E.** Fractions of spliced and unspliced reads in the three datasets. **F.** Examples of the unspliced-spliced patterns for *Nusap1*, *Ccnd3*, and *MELK* in mouse embryonic stem cells, ductal cells, and human fibroblasts, respectively.

The three datasets (mESCs, ductal cells, and fibroblasts) present different sequencing depths: mESCs and fibroblasts have ∼30 thousand unique molecular identifiers (UMI) per cell, median values of 31977 and 27319 UMIs, respectively, a depth that is uncommon for the recent standards; while the ductal cells are as low as 8 thousand UMIs per cell, median value of 8043 (Figure 1D). Similarly, the median number of genes identified per cell varies from 2840 in the ductal cells to 5161 and 5630 in fibroblasts and mESCs (Figure 1D). These differences across samples might be explained by their respective sequencing depths: total spliced and unspliced reads of 8M, 115M, and 220M in ductal cells, fibroblasts, and mESCs, respectively (Figure 1D). Overall, they have similar fractions of unspliced reads (Figure 1E). All the datasets contain genes with circular patterns in the spliced-unspliced read space in accordance with the RNA velocity theory (Figure 1F). Cycling genes are expected to be characterized by fully circular patterns as they complete both their activation and deactivation phases (Figure 2A). Overall, these datasets constitute a unique opportunity to study gene regulation throughout the cell cycle in different mouse and human cell types.

**Figure 2.**
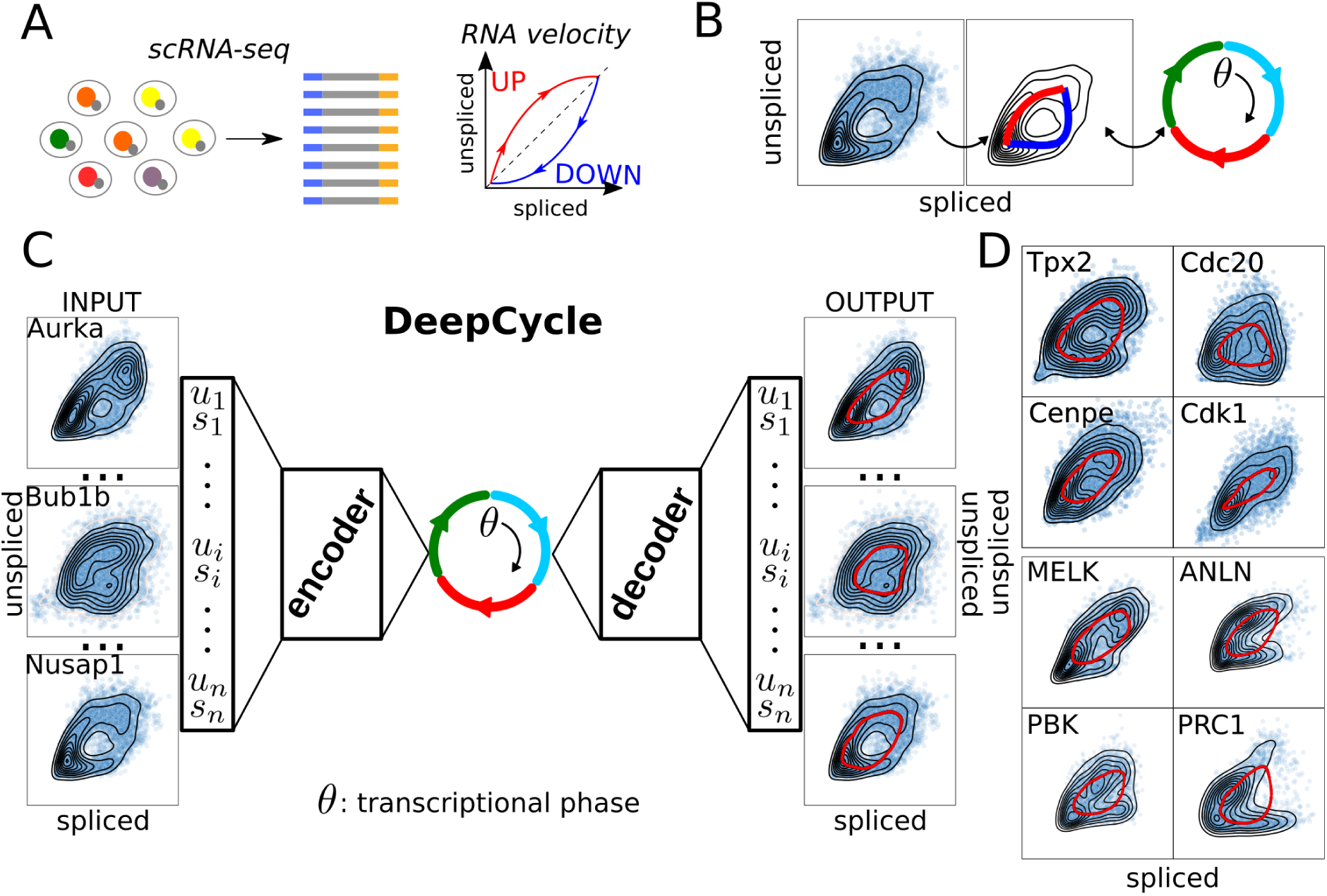
Transcriptional phase inference with DeepCycle. **A.** Single-cell RNA-seq combined with RNA velocity analysis allows the detection of transcriptional changes within a single cell. **B.** Fully circular RNA velocity patterns can be mapped to an angle describing the transcriptional state of a gene. By generalizing this to all genes, the angle will describe the actual transcriptional state of a cell. The angle is called the *transcriptional phase*. **C.** DeepCycle infers the transcriptional phase of each cell. It takes as input the spliced-unspliced reads for a set of genes. By fitting the transcriptional phase θ, it can denoise and predict the unspliced-spliced expressions for each transcriptional phase. **D.** Examples of cycling genes and fits of the RNA velocity patterns (red lines) in mESCs and human fibroblasts.

### Inference of a cell-cycle transcriptional phase from single-cell RNA-seq data

The dynamical state of a gene can be inferred by comparing its unspliced and spliced reads^9^. Unspliced reads indirectly measure the nascent transcripts, and the spliced ones the mature messenger RNAs (see Figure 1F and 2A). The comparison of the two quantities at the single-cell level allows the inference of the transcriptional activation, or deactivation, of a gene. The original framework proposed by La Manno et al.^9^ assumed either constant velocity or constant unspliced molecules; to overcome this limitation, Bergen et al.^19^ developed an extension of the original model to include intermediate states and more flexible dynamical parameters (scVelo). However, the extended model was unable to fit the actual dynamics for the genes in our datasets, while the inferred latent time did not capture the correct dynamics of the cells (see Supplementary Figure S2). Therefore, we reasoned that the complexity of gene regulation in the context of the cell cycle cannot be approximated by the current models^9, 19^ and that a more flexible approach is required. In order to achieve this, we developed a new method based on neural networks, taking advantage of their ability to represent a universal function approximator^20^.

We expect that genes whose expression is regulated during the cell cycle show a closed path in the unspliced-spliced RNA space consisting of both an active and inactive phase (see Figure 2A and 2B). Overall, the cell-cycle progression of a cell can be viewed then as a periodic trajectory within the 2N dimensional unspliced-spliced space where N is the number of considered genes. This embedded 1-dimensional manifold representing the cell cycle can be characterized by a circular latent variable, the *transcriptional phase* (θ), that maps cells into the particular location of the periodic trajectory. Notice that θ is a continuous variable representing the continuous cell-cycle progression of cells that has not to be confused with the discrete phases of the cell cycle (G1, S, G2, and M). Then, the estimation of θ for each cell given the unspliced and spliced reads is an embedded manifold learning problem. To solve this problem, we developed DeepCycle, a deep learning method based on an AutoEncoder (AE) neural network. AEs are designed to perform non-linear dimensionality reduction by compressing the information contained in the inputs to a lower-dimensional space (latent space) in the encoding phase. The compressed information is then used to reconstruct the original input in the decoding phase. AEs have been used to analyze scRNA-seq data and accomplish different tasks, from clustering to denoising^21–28^. DeepCycle is constructed as an AE with a single latent variable representing the cell-cycle transcriptional phase θ that is then transformed with cosine and sine functions in the first layer of the decoder (Figure 2C and Supplementary Figure S3).

To train DeepCycle, we used the expression of unspliced and spliced RNAs of the genes in the GOterm:cell_cycle (n=532, see Methods) determining circular paths for cycling genes in the unspliced-spliced space and removing technical noise or biological fluctuations associated with stochastic gene expression (see examples in Figure 2C-D and Supplementary Figure S4). Finally, a transcriptional phase is assigned to each cell in the dataset (see Methods) and the dynamics of unspliced and spliced RNA with respect to the transcriptional phase can be further analyzed. It is important to note that the transcriptional phase is a non-linear monotonic function of time that can be arbitrarily complex, so we cannot directly infer temporal dynamics with it. Importantly, DeepCycle robustly returns very similar transcriptional phases by selecting as input the genes showing multiple maxima in the unspliced-spliced space (Supplementary Figure S5 and Methods). The genes presenting multiple maxima (n=158) are listed in Supplementary Table S1 that includes cycling genes not yet considered in the GO term:cell_cycle that could be added as markers of the cell cycle.

Finally, we compared DeepCycle with Cyclum^7^, a recent method developed for the analysis of the cell cycle in scRNA-seq data also based on an AE. Strikingly, Cyclum was not able to place cells consistently in a circular 1D manifold and therefore could not correctly identify the cell-cycle progression of single cells when applied to our datasets (Supplementary Figure S6). As opposed to Cyclum, DeepCycle is based on RNA velocity and trained on both spliced and unspliced RNA levels which may explain the better performance. Moreover, DeepCycle produces dynamic trajectories in the unspliced-spliced space for each gene (see Figure 2D) and the quality of the fit of each trajectory to the data can be used to evaluate whether the learning process worked properly.

### Detection of cell cycle phases in multiple cellular models

Single cells can be associated with the S and G2/M phases by analyzing the expression of representative marker genes^19, 29^. For example, the cell cycle scores calculated by scVelo match well with the transcriptional phases inferred by DeepCycle (Supplementary Figure S7). To better define the transitions between cell cycle phases we looked at the expression of specific marker genes (Figure 3A). The G1/S transition corresponds to the peak in cyclin-E2, the S/G2 transition to the increase of the mitotic cyclin-F, and mitosis to the sharp decrease of *Wee1*/*WEE1*. The loss of *Wee1/WEE1,* a protein kinase inhibiting mitosis, allows the cyclin B1-Cdk1 complex to activate the cascade of reactions necessary to proceed into mitosis^30, 31^ and, consistently, the mRNA levels of the Aurora kinases A (*Aurka*/*AURKA*) that localize at the centrosomes^32, 33^, and of the Nucleolar and spindle associated protein 1 (*Nusap1*/*NUSAP1*), that plays a role in spindle microtubule organization^34, 35^, increase in G2 and M phases (Figure 3B). Other possible marker genes show cycling patterns as expected, e.g. *Orc1*/*ORC1*, *Mcm6*/*MCM6*, *Ccne1*/*CCNE1*, *Ccna2*/*CCNA2*, and *Ccnb2*/*CCNB2* (Figure 3A).

**Figure 3.**
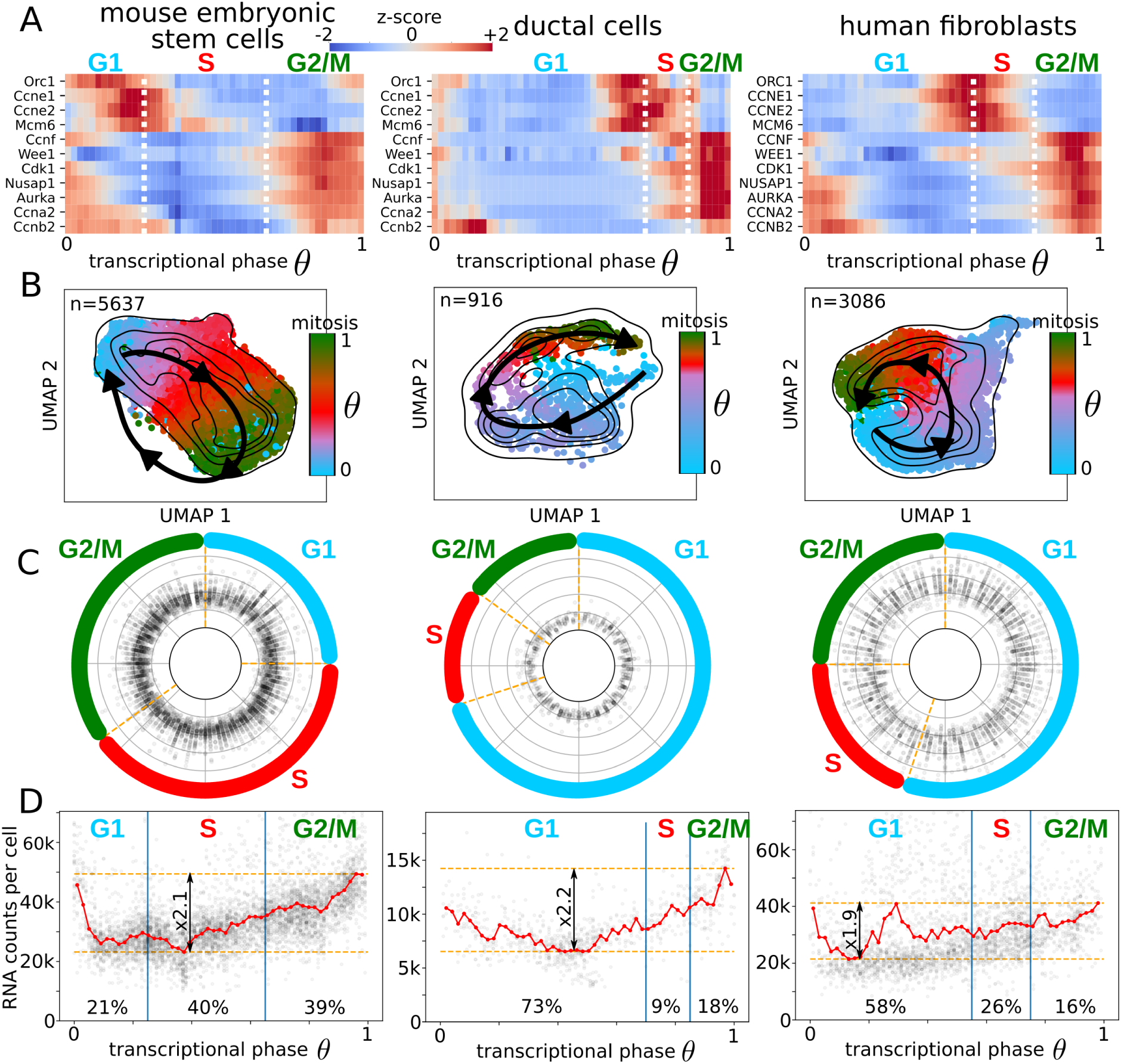
Cell cycle analysis in mouse and human cellular models. **A.** The transcriptional phases and the cell cycle phases were connected based on marker genes. Z-scores are only intended for comparison purposes, the changes in the expression across the cycle are, in general, highly significant. **B.** The UMAP embeddings for mESCs, ductal cells, and human fibroblasts with the cell cycle directionality identified by DeepCycle (black arrows). The cells are associated with different colors depending on the cell cycle phase they belong to, light blue for cells in G1, red in S, and dark green in G2/M. **C.** The transcriptome variabilities (χ^2^) are constant across the transcriptional phases. **D.** RNA counts per cell as a function of the transcriptional phase show doubling trends followed by a sudden drop that identifies mitosis. The fibroblasts show contamination of cells from the nonproliferative subpopulation, arrested in mid-G1, as discussed in the last section.

To simplify the comparison of the cell cycles across datasets, the transcriptional phases were normalized between 0 and 1 and were aligned such that mitosis occurs approximately at θ=1. The paths of the cells around the cell cycle can easily be identified in the 2-dimensional projections (Figure 3B) and the variability of the transcriptomes across the transcriptional phase is stable (Figure 3C). Though the extended RNA velocity model^19^ did not capture the correct dynamics at the level of the single gene (Supplementary Figure S2), it could infer the correct dynamics of transcriptional changes at the cell level (see the velocity plots in Supplementary Figure S8).

Fast cell cycles are typically associated with pluripotency and stemness^11–13^. Consistently, the mESCs present the lowest number of cells in G1, while fibroblasts and ductal cells have much more extended G1 phases (see Figure 3D). The fractions of mESCs assigned to the different phases are 21% to G1, 40% to S, and 39% to G2/M (Figure 3D). Similar fractions are identified by flow cytometry analysis, respectively, 22-26% in G1, 42-51% in S, and 27-32% in G2/M (Supplementary Figure S9).

At mitosis, the mother cell needs to have approximately double its original volume in order to generate two daughters of the same initial size. Droplet-based single-cell technologies, such as 10x, can indirectly detect the different cell sizes, where a bigger cell means a higher concentration of mRNA within the droplet, which should reflect a higher count of unique RNA molecules (UMIs) within the cell. In this case, the increase in the unique RNA molecules across the cell cycle should be roughly proportional to 2. Indeed, as predicted, the RNA counts per cell as a function of the transcriptional phase show a positive fold change of 2.1, 2.2, and 1.9 for mESCs, ductal cells, and fibroblasts, respectively (Figure 3D). Further, the flow cytometry analysis performed for the mESCs showed roughly a doubling size passing from G1 to G2/M phases (Supplementary Figure S9). After validating that the transcriptional phases identified by DeepCycle are consistent with the global features of the cell cycle, such as cell cycle markers, cell sizes, and fractions of cells in each phase, we can discuss the regulation of individual cell cycle genes at the mRNA level.

Members of the Cdc25 family are well conserved key regulators of the cell cycle^36–38^. The mRNA expression of the Cdc25 family of proteins shares the same behaviour across the datasets, i.e. Cdc25a/CDC25A increases at the G1/S transition while Cdc25b-c/CDC25B-C at the G2/M (Supplementary Figure S10), consistently with the function of their protein products^39^.

The minichromosome maintenance protein complex (Mcm) is a heterohexamer, formed by Mcm2-7/MCM2-7 proteins, which works as a helicase that unwinds the double-stranded DNA and powers the replication fork progression during the S phase^40^. As expected, the mRNA levels of all subunits of the Mcm peak at the beginning of the S phase for all the datasets (Supplementary Figure S10). Interestingly, other DNA replication genes, such as components of the Origin recognition complex proteins (Orc1-6/ORC1-6), show different expression patterns across the datasets, suggesting more heterogeneous regulation (Supplementary Figure S10).

*Cdk1/CDK1* mRNA level increases in G2 and M phases as required by its protein function^41^ (Figure 3A). The other main Cdk mRNAs (*Cdk2-4-6/CDK2-4-6*) show lower expression levels across phases and are less consistent across the datasets, they might rather be regulated at the protein level, translationally or posttranslationally (Supplementary Figure S10). It has been previously shown that protein levels of the cyclin-E and A do not change across the mESC cell cycle^42^, but instead, mRNA levels are upregulated at the G1/S transition and in the G2/M phase, respectively (Figure 3A).

Finally, DeepCycle allows a genome-wide investigation of gene expression dynamics across the cell cycle. Indeed, we observed different waves of gene expression during the different phases of the cell cycle (Supplementary Figure S11). Interestingly, a similar fraction of genes reaches their maximum RNA level in each phase in mES cells. On the contrary, in ductal cells and human fibroblasts, most genes reach the maximum level in G1. Overall, DeepCycle consistently identifies cycling genes and shows their mRNA synthesis rate (unspliced) and expression level (spliced) across the cell cycle.

### Identification of cell-cycle core transcription factors

Having characterized the cell cycle at the mRNA level, another feature of our approach is that it allows us to identify the potential transcription factors (TFs) responsible for the gene expression dynamics. Transcription factors bind to DNA-specific sequences (binding motifs) and activate the transcription of their target genes. They encode the cellular programs for many of the functions a cell needs to perform. To infer the TFs active during the cell cycle, we implemented an ISMARA-like approach^43^. Briefly, Balwier et al. introduced a linear model to infer TF activities from bulk RNA-seq samples. To apply it to our data, we used the same linear model to try to explain the expression level of the unspliced reads in single cells. Even if the amount of unspliced reads is much lower compared to the spliced reads (∼5-6 times less, Figure 1E), they remove the effect of mRNA stability, reflecting more closely the nascent transcription events and, therefore, the effect of transcription factors at the gene promoters.

The motif analysis shows that most of the TF activity takes place in the G1 phase when cells need to decide whether to go into another round of replication or to arrest the cycle in order to accomplish a new function (Figure 4A). Among the most significant TFs, Yy1/YY1 has been identified in all the datasets as active in G1 phase, suggesting a general role during the cell cycle (Figure 4A-B). Indeed, Yy1 is known to induce proliferation and maintain pluripotency of mESCs through the BAF complex^44^. Interestingly, Yy1 binds to chromosomes during mitosis^45^ and, accordingly, its transcription starts already in the G2/M phase suggesting a pioneering activity at the beginning of a new cycle (Figure 4B).

**Figure 4.**
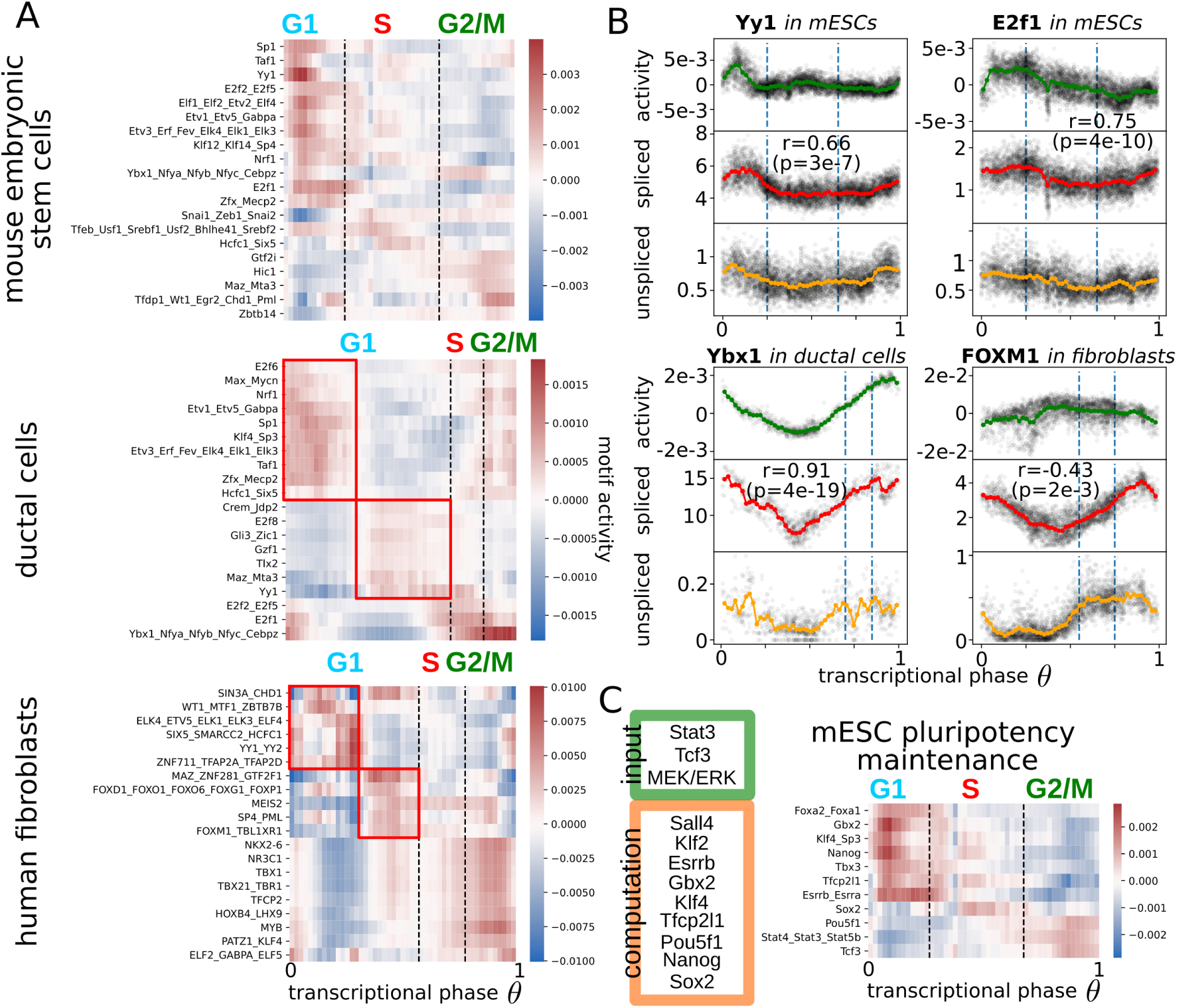
Transcription factor dynamics driving expression during the cell cycle. **A.** Motif activities for families of transcription factors across the cell cycle in mESCs, ductal cells, and human fibroblasts. The red boxes identify two waves of transcription in the G1 phases for more differentiated cell lines. **B.** The comparison of motif activities with the respective mRNA levels around the cell cycle for Yy1 and E2f1 in mESCs, Ybx1 for ductal cells, and FOXM1 for human fibroblasts. These comparisons allow clarifying whether the regulation is happening more at the transcriptional level or the protein level. **C.** The two sets of TFs in the green and orange boxes are at the core of the pluripotency maintenance network in mESCs^57^. The heatmap shows the activities of the pluripotency factors around the cell cycle of mESCs.

For both mouse datasets (mESCs and ductal cells), the E2f family appears as a critical group of regulators. Members of the family are known to act at the beginning of the cell cycle specifically for the G1/S transition and to become active after the phosphorylation of the retinoblastoma proteins (pRb)^1, 46, 47^. Two E2f-related motif activities (E2f1, E2f2_E2f5) peak in between the G1 and the S phase, presumably to activate the genes necessary for the transition^48^ (Figure 4A). More specifically in mESCs, E2f1 seems to be mostly regulated at the protein level since the change in the mRNA level (∼50%) is very little compared to the change in activity (Figure 4B). Other factors seem to act similarly between mESCs and ductal cells, like the TATA-binding protein-associated factor (Taf1), the Specificity factor 1 (Sp1), and the Nuclear respiratory factor 1 (Nrf1), all active in early G1 (Figure 4A).

Regarding the ductal cells, we found a very high correlation (r=0.91, p=4e-19) between Ybx1 mRNA level and the activity of its motif where both are constantly increasing from G1 to M (Figure 4B). Interestingly, Ybx1 positively regulates the G1 and G2/M phases of the cell cycle^49, 50^ and its expression is linked to poor prognosis in pancreatic ductal adenocarcinoma^51^. Regarding the factors appearing in the human fibroblasts, MYB plays a role in the G2/M transition, with a constant increase of expression from G1 to G2/M^52, 53^. Also, its targets follow the same trend, and MAZ induces MYB expression shortly after the exit from quiescence, bypassing the inhibition of E2F-pRB^54^ (Figure 4A). Similar to Ybx1 in ductal cells, the mRNA level of FOXM1 grows constantly from G1 to M as expected by its function during mitosis^55, 56^, but the activity of its targets is slightly anticorrelated, hinting at a complex post-transcriptional regulation^56^.

For mESCs the maintenance of the pluripotent state is crucial and the main factors involved in the pluripotency transcriptional program are known^57^ (Figure 4C). Among them, the strongest activation happens for the targets of Stat3^58^/Stat4/Stat5b, Tcf3, and Pou5f1 (Oct4), which are increased in G2/M, followed by Klf4^59^/Sp3, Gbx2, Nanog, Tfcp2l1, and Essrb^60, 61^/Essra in G1 (Figure 4C).

From a general perspective, a clear pattern emerges by comparing the undifferentiated mESCs with the more differentiated human fibroblasts and ductal cells. The undifferentiated cells show a strong and unique wave of activation of TFs in G1. Instead, in the more differentiated cell types, the activities of the TFs across the G1 phase cluster into two groups. The first group displays an early activation directly after mitosis, while the second group exhibits a late G1 activation (red boxes in Figure 4A). We believe these waves are linked to cell-fate decisions, as discussed in the next section.

### Characterization of cycling cells shifting to the cycle-arrested state

The human fibroblasts include a subpopulation with a low cycling activity, that we excluded from the previous analysis (Supplementary Figure S1). By mapping the ‘nonproliferative’ cells across the cycle with the model trained with DeepCycle, this sub-population was associated with the mid-G1 phase (Figures 5A-B). Therefore, the full population of fibroblasts comprises 76% in G1/G0, 15% in S, and 9% in G2/M phases (Figure 5B). Similar numbers have been obtained through flow cytometry analysis, where the DNA content assigns 79% to G1/G0, 13% to S, and 8% to G2/M phases (Figure 5C). Further, the flow cytometry analysis shows that some cells in G1 are bigger than the cells in S and G2/M suggesting that the cells waiting in G0/G1 are increasing in size, it remains unclear whether they will re-enter the cycle later. The cell velocities are consistent with our interpretation and do not clarify if the cells in G0 will start cycling again (Supplementary Figure S12).

**Figure 5.**
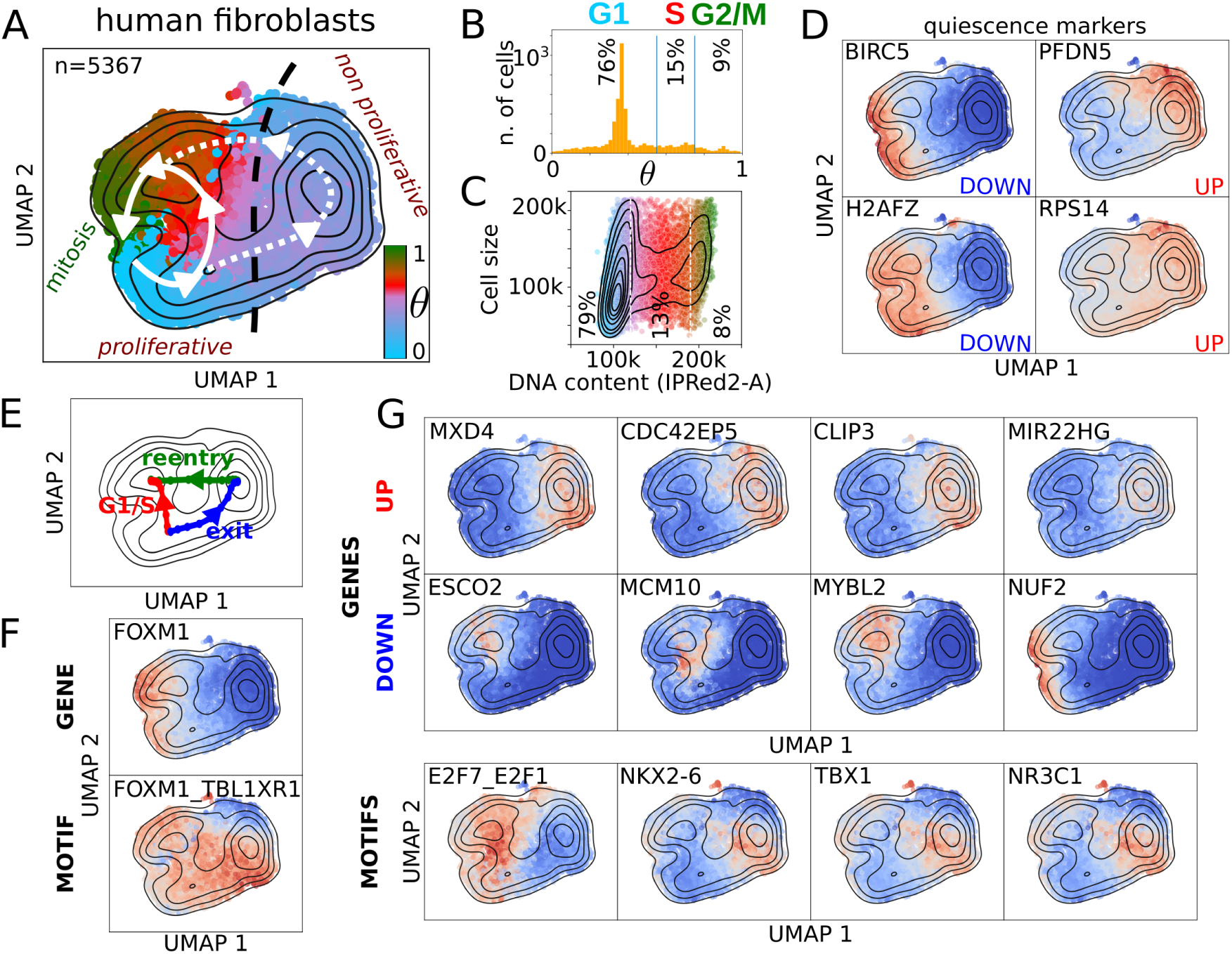
G1-to-S and G1-to-G0 transitions in human fibroblasts. **A.** The two subpopulations of human fibroblasts, separated by the black dashed line: on the left side the proliferative cells, and on the right the nonproliferative. Each cell is colored according to its inferred transcriptional phase. **B.** The distribution of transcriptional phases for the human fibroblasts shows the nonproliferative fibroblasts arrested around θ=0.4. **C.** Flow cytometry analysis recapitulates fractions of cells in the main phases of the cell cycle similar to the phases identified with DeepCycle. The cells in G0/G1 can be bigger than the cells in G2/M. The colors are consistent with panel A, G1 in blue, S in red, and G2M in green. **D.** Examples of G0 marker expressions for the two subpopulations^62^. BIRC5 is also known as survivin. **E.** The paths, going from mid-G1 towards the S phase (in red), from the mid-G1 phase towards the nonproliferative state (in blue), and the nonproliferative state towards the S phase (in green), are identified by the higher density of cells. **F.** FOXM1 is downregulated in the nonproliferative cells and shows the opposite trend along the G1/S and exit paths (red and blue in panel E). The activity of its motif is completely uncorrelated with its expression. **G.** The top up- and down-regulated genes with the strongest fold changes comparing the paths in E. The third row shows the most significant motif activities identified by comparing the paths in E.

The two subpopulations split their trajectories around mid-G1 (Figure 5A) and specific markers of quiescence^62^ suggest the nonproliferative cluster might be entering into the G0 phase (Figure 5D). To detect the underlying changes in the gene expressions and their regulations, we implemented a method based on a modified version of the Nudged Elastic Band^63^, to infer the paths connecting the cell states in the bidimensional space (Figure 5E). The detected paths follow the trajectories with the highest density of cells, as shown in Figures 5E.

To strengthen our hypothesis of the nonproliferative cells being quiescent, we checked the expression along the two paths of G0 markers^62^. The markers supposed to be G0-downregulated are consistently inactivated (Supplementary Figure S13) while the signal for the G0-upregulated genes is unclear (Supplementary Figure S14). Therefore, nonproliferative fibroblasts might represent a differentiated state of the fibroblasts and not simply reflect cells entering in G0^64^. FOXM1 is strongly downregulated in nonproliferative cells, making it a good candidate as a marker of quiescence and one of its potential regulators. With regards to the cell cycle analysis (Figure 4B), FOXM1 targets do not follow its mRNA expression, but are upregulated while exiting the cell cycle (Figure 5F). Among the top genes upregulated along the path towards the quiescent state (the blue and green paths vs the red path in Figure 5E), we find MXD4, CDC42EP5, CLIP3, and MIR22HG. MXD4 is a MYC antagonist known to increase the fraction of cells in the G0/G1 phase in hematopoietic differentiation^65^, and could be a master regulator of entry into the G0 phase. CDC42EP5 is a small Rho-GTPase belonging to the Borg family and is involved in cell shape regulation and lamellipodia formation^66^. Similarly, CLIP3 (or CLIPR-59) is a CAP-Gly domain containing linker protein with a poorly-specified function, perhaps modulating the compartmentalization of the AKT kinase family^67^. Lastly, MIR22HG is a long non-coding RNA involved in proliferation that acts as a tumor suppressor in primary lung tumors ^68^ and leads to poor prognosis in glioblastoma^69^. On the other side among the most downregulated genes in the quiescent state are ESCO2, MCM10, MYBL2, and NUF2. ESCO2 is needed during the S phase to modify cohesin^70^ and MCM10 accumulates during the S phase, while being lowly expressed during the rest of the cycle^71^. MYBL2 (B-Myb) belongs to the family of the MYB transcription factors and has been typically associated with poor prognosis in cancer^72^, while NUF2 localizes at centrosomes and is necessary for mitotic progression in vertebrates^73, 74^. For the extended lists of up and down-regulated genes see Supplementary Figures S15-S18.

Importantly, the top motif, that distinguishes the two subpopulations (E2F7_E2F1), belongs to the E2F family, which is one of the master regulators of the cell cycle^1^, and is strongly inactivated in the non-proliferating quiescent cells. The other TFs shown in Figure 5G (NKX2-6, TBX1, and NR3C1) do not have a clear function associated with the cell cycle, so further studies are needed to elucidate their role. More TF motifs associated with the paths are shown in the Supplementary Figures S19-S20.

In summary, DeepCycle allowed us to characterize the G1-G0 transition in wild type fibroblasts without having to perturb the cells, finding novel candidate genes and transcription factors regulating quiescence.

## Discussion

We generated scRNA-seq datasets in mouse embryonic stem cells and human fibroblasts with high sequencing depth. The circular RNA velocity patterns emerged clearly in cell-cycle regulated genes revealing the activation/inactivation phases that these genes undergo during the cell cycle. We developed DeepCycle, a novel deep learning approach, to exploit the RNA velocity patterns and study gene regulation dynamics during the cell cycle. DeepCycle assigns a cell-cycle transcriptional phase for each cell by fitting the RNA velocity patterns. Furthermore, the inferred transcriptional phase can be associated with cell-cycle phases thanks to known gene markers. Thus, DeepCycle allows us to determine the cell-cycle progression state of each cell from scRNA-seq data and identify novel genes involved in the cell cycle. Importantly, the efficacy of the method was extensively proven in cellular models from different organisms at different developmental stages.

The decision to implement a novel approach came after noticing the failure of the current methods within the RNA velocity framework^7, 19^, to correctly infer the dynamics of the cycling genes. DeepCycle’s ability to infer cycling patterns in the spliced-unspliced RNA space at the gene level shows that the framework of the RNA velocity can be further improved by the study of more flexible models of transcription. Likely the assumptions in the previous model (constant rates) should be relaxed to fit the transcriptional model to the data. It is reasonable to imagine that the transcription, splicing, and degradation rates are complex functions changing during the cell cycle progression. Our method will allow the analysis of trajectories without making assumptions about the model parameters, enabling more focus on the dynamics of the single gene.

The analysis highlighted known and novel cell cycle regulators in established cell lines, identifying two major waves of transcription in the G1 phase of differentiated cells while pluripotent cells seem to undergo a single wave of transcription during G1. The two waves are likely to be associated with the restriction point where the cells finally commit to undergoing another cell cycle. Further, for the first time, we could observe single cells exiting from the cell cycle in an scRNA-seq sample and disentangle the underlying regulations, thereby providing lists of novel targets for the regulation of the cell cycle and the quiescent states in mammalian cells. We envision that our approach will facilitate the characterization of the branching point between the S and G0 phases in multiple cellular models by applying it to other scRNA-seq datasets. In particular, an extensive study of the transcriptional changes happening at the cell cycle while cells reach confluence is still missing and of general interest.

Finally, we anticipate that DeepCycle will become an essential tool for the scientific community to further investigate the cell cycle in a broad range of systems without the need for cell synchronization or genetic-tagging. This makes our approach especially suitable to study the interplay of the cell cycle with pluripotency and cell reprogramming^75^. Moreover, the comparison between normal and cancer tissues may lead to the discovery of cell-cycle dysregulated mechanisms in tumors and, perhaps, potential targets for drug development.

## Methods

### Cell culture

E14Tg2a.4 mouse embryonic stem cells were cultured on 0.1% gelatin-coated culture plates in DMEM (4,5g/l glucose) supplemented with GLUTAMAX-I, 15% heat-inactivated fetal calf serum (42F5874K, ESC culture tested, GIBCO), 0.1 mM beta-mercaptoethanol, 0.1 mM non-essential amino acids 1,500 U/ml leukemia inhibitory factor (produced in house), 3 µM CHIR99021 (72054, Stem Cell Technologies) and 1 µM PD0325901 (72184, Stem Cell Technologies) in 5% CO_2_ at 37°C.

IMR90 primary human fetal lung fibroblast cells were cultured in DMEM 41966 (4,5g/l glucose) supplemented with 10% fetal calf serum, Penicillin 100 UI/ml, and Streptomycin 100 µg/ml in 5% CO2 at 37°C. The cells were at passage 21 when performing the experiments. For both scRNAseq and FACS experiments, 20 000 cells per well were seeded into 6 well plates and cultured for 72 hours.

### Single-cell RNA sequencing

To obtain single cell suspension of mESCs for single cell RNA sequencing, cells on a 60 mm culture dish were washed once with PBS and treated with 1 ml 0,25% trypsin-1mM EDTA (25200-072, Invitrogen) at 37°C for 3 minutes, then harvested into 3ml medium containing serum, and washed 2-times with PBS containing 0.04% BSA. To prepare single cell suspension of IMR90 cells, the cells from one well of a 6 well culture plate were washed twice with PBS and treated with 500ul 0.05% trypsin-0.53mM EDTA (25300-062, Invitrogen) at 37°C for 2 minutes, then harvested into 4.5ml medium containing serum, passed through 50µm cell strainer and washed 2-times with PBS containing 0.04% BSA. In both cases, cell concentration and viability (98 %) was determined using Countess II (Invitrogen) according to the manufacturer’s instructions. Cells were then processed using the 10x Genomics Chromium System according to the manufacturer’s instructions.

Cell number and viability were determined by a Trypan Blue exclusion assay on a Neubauer Chamber. Samples consisting of > 90 percent viable cells were processed on the Chromium Controller from 10x Genomics (Leiden, The Netherlands). Ten thousand cells were loaded per well to yield approximately 6500 captured cells into nanoliter-scale Gel Beads-in-Emulsion (GEMs).

In the case of mESCs, the single-cell 3 prime mRNA seq library was generated according to 10X Genomics User Guide Chromium Single Cell 3ʹ Reagent Kits v3 (P/N CG000183 Rev A). For the human fibroblasts, the single-cell 3 prime mRNA seq library was generated according to 10x Genomics User Guide Chromium NEXT GEM Single Cell 3’ Reagent Kits v3.1 (P/N CG000204 Rev D). The raw and processed data for both libraries were stored on the GEO (Accession number: GSE167609).

CellRanger outputs have been processed with velocyto^9^ (version 0.17.17) and analyzed using scanpy^76^ (version 1.4.4.post1) and scvelo^19^ (version 0.2.2).

### Cell cycle assay and flow cytometry

Cells were harvested by trypsin as before and washed once with PBS. About 2x10^6^ cells were resuspended in 100 µl PBS and added drop-by-drop to 900 µl 95 % ethanol while mixing, then stored at +4°C overnight. Cells were then collected by centrifugation, washed once with PBS, re-suspended in 1 ml staining buffer (50 µg/ml propidium iodide, 2 mM MgCl_2_, 50 ng/ml RNaseA [EN0531, ThermoScientific] in PBS) and incubated for 20 minutes at 37°C. Stained cells were washed once with PBS and analyzed on BD LSRII flow cytometer.

The fcs files were processed with fcsparser (https://github.com/eyurtsev/fcsparser). For the mESCs, the debris in the data was removed by filtering SSC-H and SSC-W values higher than 140000 and 100000, respectively, and by selecting cells with a Hotelling T^2^ value lower than 6 in the FSC-A SSC-A space, see Supplementary Figure S8. The filtering retained more than 80% of the original cells (∼ 26k out of 31k). For the human fibroblasts, SSC-H values lower than 25000 and, SSC-H and SSC-W values greater than 150000 and 110000, respectively, were excluded. As for the mESCs, only cells with Hotelling T^2^ lower than 6 in the FSC-A SSC-A space were retained.

### Implementation of DeepCycle

The autoencoder was implemented in TensorFlow 2. The training happens in two steps, first, we train the encoder to predict the cell angles in respect to the average spliced-unspliced of a selected gene (Nusap1 for mESCs, Ccnd3 for ductal cells, and MELK for human fibroblasts). Second step training of the whole autoencoder on unspliced-spliced data of the genes in the GOterm:cell_cycle (GO:0007049). The structures of the encoder and decoder are depicted in Supplementary Figure S3. The Dense layers are activated with a Leaky ReLu function and the transcriptional phase θ in the hidden dimension is mapped to (Cos(θ), Sin(θ)). During the training Gaussian Noise was added before outputting the θ from the encoder at (Cos(θ), Sin(θ)) in the decoder.

Finally, to infer a phase for each cell, we binned the angles in 50 and assign a cell to the closest bin in the unspliced-spliced space predicted by the autoencoder (red lines in Figure 2C-D and Supplementary Figure S4) for all the genes used for the training (GO term: cell_cycle, GO:0007049).

DeepCycle implementation was stored in the GitHub repository https://github.com/andreariba/DeepCycle.

### Transcription factor activity

The linear model used to infer the motif activities was implemented as in ISMARA^43^. To find the regulatory interactions between transcription factors and genes, we used Motevo predictions of binding sites in promoters downloaded from the Swiss Regulon Portal (https://swissregulon.unibas.ch/sr/downloads) for mm10 mouse genome assembly (https://swissregulon.unibas.ch/data/mm10_f5/mm10_sites_v2.gff.gz) and hg19 human genome assembly (https://swissregulon.unibas.ch/data/hg19_f5/hg19_sites_v2.gff.gz). Briefly, the Motevo algorithm uses a Bayesian framework to estimate the posterior probability that a binding site for a given weight matrix (associated with a motif) occurs in an interval^77^. After, we summarized the transcription factor binding sites in a matrix of site-counts N_pm_ by summing the posterior probabilities for each motif *m* in a promoter *p*. We defined a promoter as the TSS +/- 1kb.

The cross-validation was repeated 10 times and the average optimal strength of the ridge regularization was used for the final calculation of the TF activities.

### Identification of cycling genes and high-density paths

A mixture of two bivariate Gaussians was used to fit the distribution of unspliced-spliced expressions, to identify genes showing at least two maxima in the distribution of cells. After identifying the two Gaussians a Hotelling’s T^2^ test was applied to select the genes with two significantly different attractors. After supervised filtering of the remaining genes, we implemented a method to select genes with at least two paths connecting the two maxima. The path detection was implemented into two steps. First, coarse-grained paths were drawn by slicing the spliced-unspliced landscape and connecting the minima found across the successive slices. To refine the identified paths we implemented the Nudged Elastic Band^63^ with the addition of a viscosity term to stabilize the dynamics, we called the method Viscous Nudged Elastic Band (VNEB).

The VNEB was also applied to the fibroblasts dataset to identify the paths connecting the G1 phase to the S and G0 phases in Figure 5E.

## Acknowledgments

We thank the cell culture service, the flow cytometry service and the imaging center of the Institut de Génétique et de Biologie Moléculaire et Cellulaire (IGBMC). We thank specially the national platform GenomEast for the sequencing and the 10x Genomics reagents.

This study was supported by funds from Conseil National de la Recherche Scientifique, Institut National de la Santé et de la Rechrche Médicale and Université de Strasbourg; the grant ANR-10-LABX-0030-INRT, a French State fund managed by the Agence Nationale de la Recherche under the frame program Investissements d’Avenir ANR-10-IDEX-0002-02; and, the fellowship «IDEX chaires attractivité recherche» granted by the University of Strasbourg. Work in the lab of WMK was funded in part by grants from: La Fondation Recherche Medicale (FRM) (AJE20160635985), La Fondation Schlumberger pour l’Education et la Recherche, FSER 19 (Year 2018)/FRM and La Ligue Contre le Cancer.

## Author contributions

AR, AO, NM designed the research. AR and NM designed the method. AR implemented the method. AO, MD, VA, and MC performed wet-lab experiments to generate the sequencing data. AR, AO, and MD performed the flow cytometry analysis. AR, MJ, and CK performed the bioinformatic analysis of the sequencing data. AR and SJC performed the motif analysis. AR and NM wrote the paper. AO, MJ, SJC and WMK critically read and edited the paper. NM and WMK conducted supervision and obtained grants to fund the research.

## Supplementary Figures

**Supplementary Figure S1.**
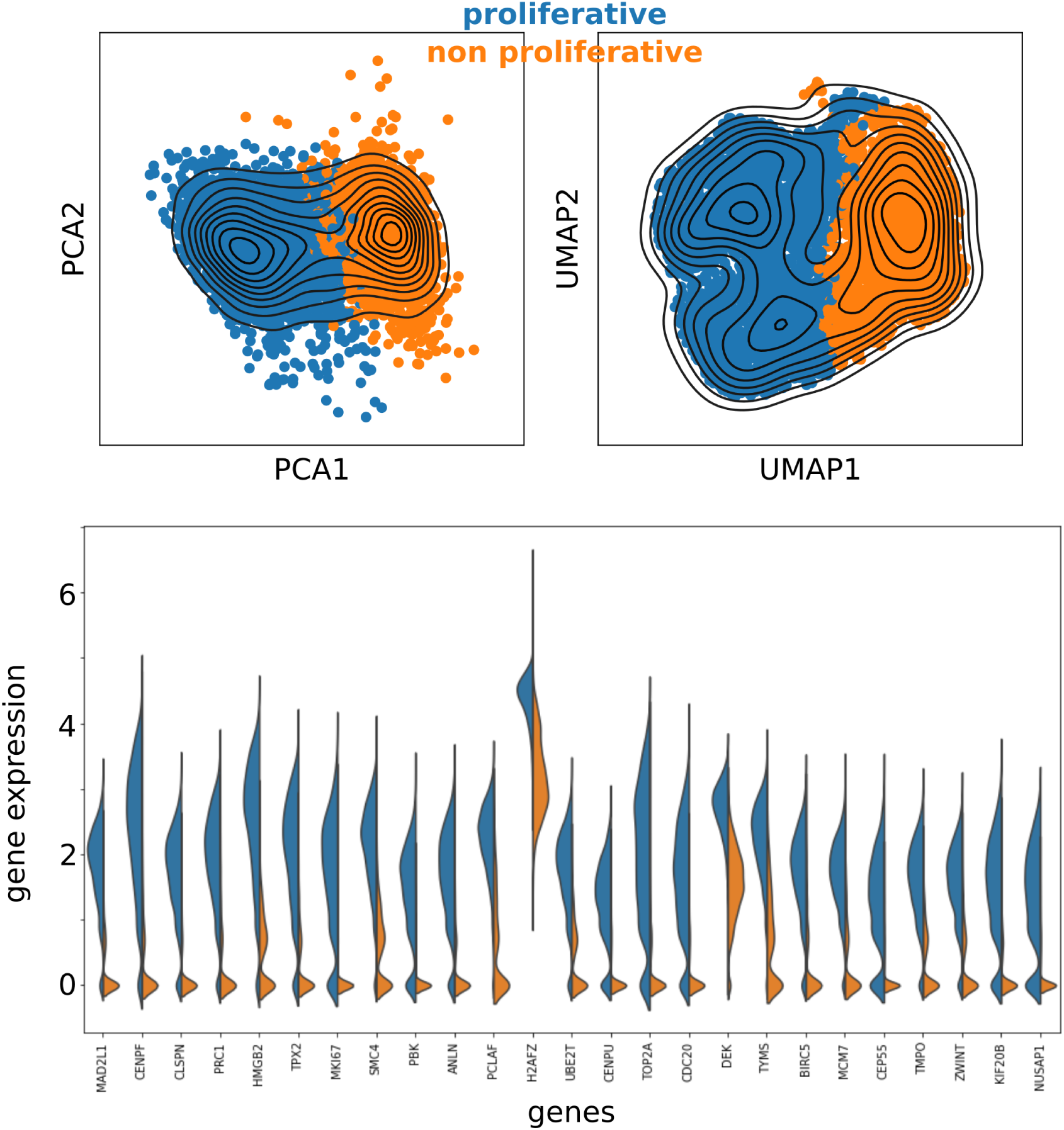
The human fibroblasts dataset contains two subpopulations, one expressing cell cycle genes (blue) and the other not expressing them (orange). The two populations are distinguishable in the PCA and UMAP projections. Leiden clustering was performed to assign cells to the two subpopulations. The top genes identifying the proliferative cluster against the nonproliferative ones are mostly associated with mitosis (DAVID UP_Keywords Benjamini=1e-13) and cell cycle (DAVID UP_Keywords Benjamini=2e-15).

**Supplementary Figure S2.**
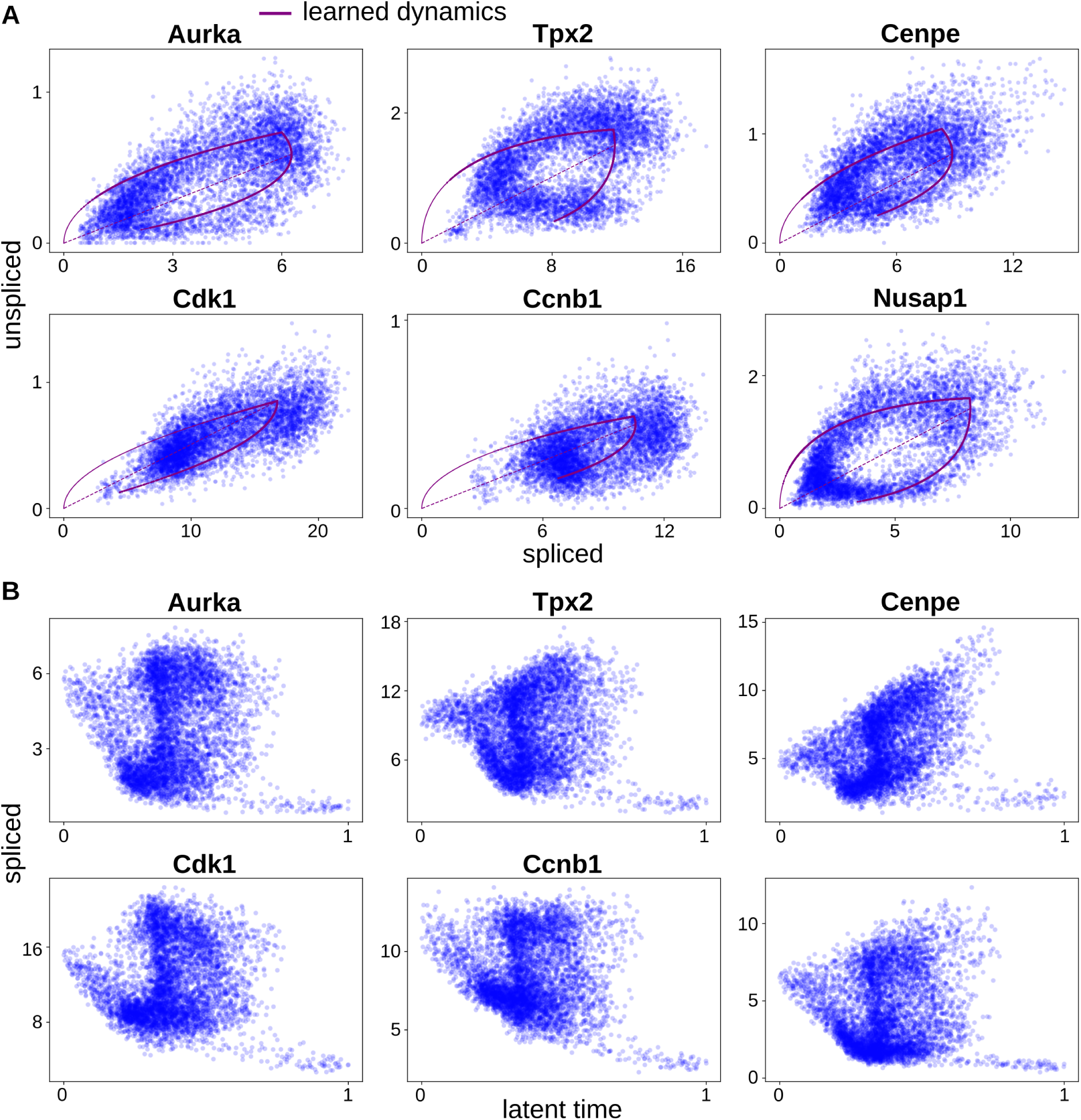
**Generalized RNA velocity (scVelo**^19^**) cannot fit the correct model and latent time**. **A.** Examples of models learned by scVelo that do not fit correctly the data. The main issue seems to be related to the inability to capture the lower steady-state. **B.** The expression of the genes as a function of the latent time shows inconsistent patterns.

**Supplementary Figure S3.**
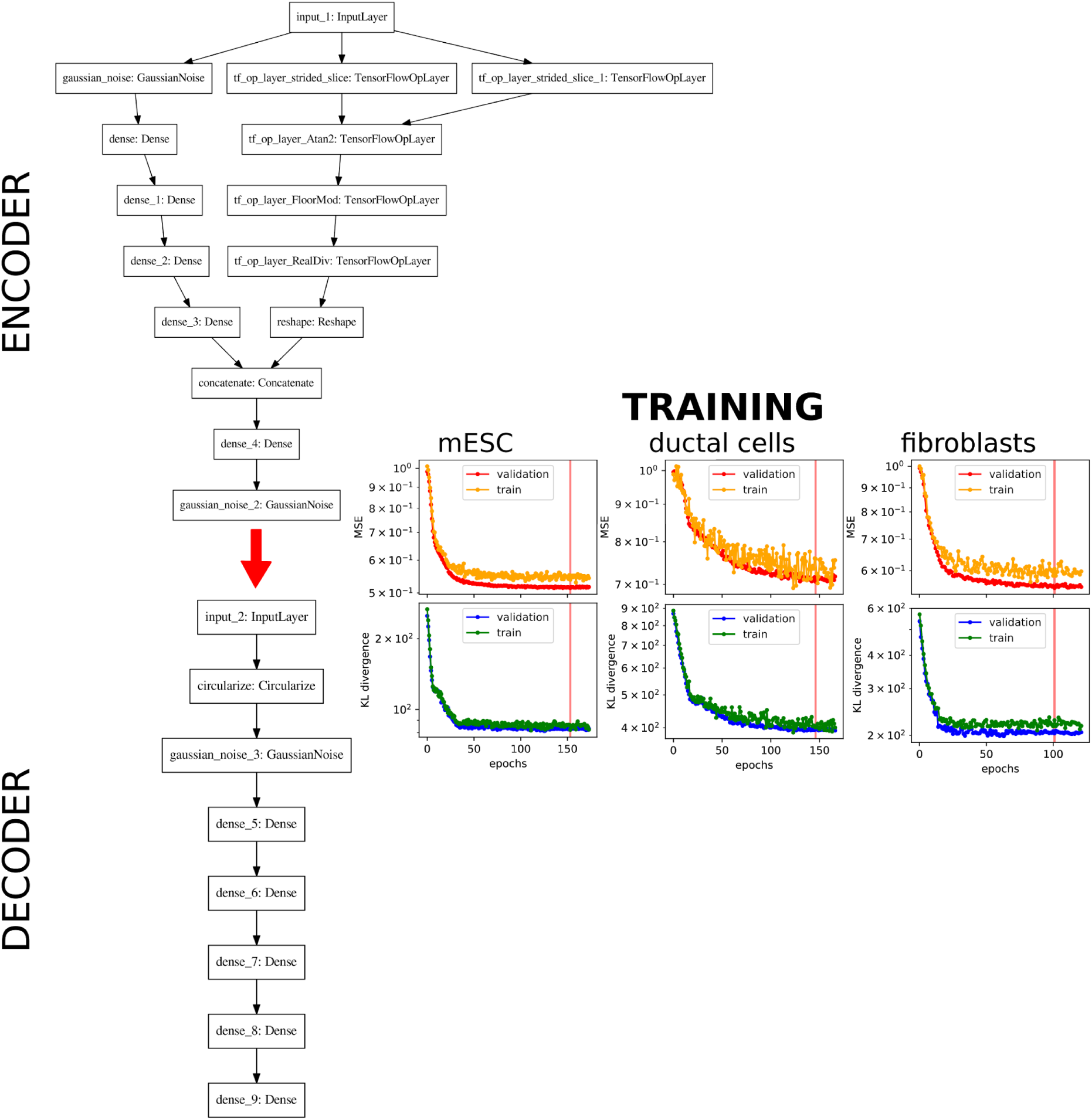
**Autoencoder structure and training**. Encode and Decoder: key layers are the Circularize, which transforms the angle with cosine and sine, and the GaussianNoise, which adds white noise to the transformed values.

**Supplementary Figure S4.**
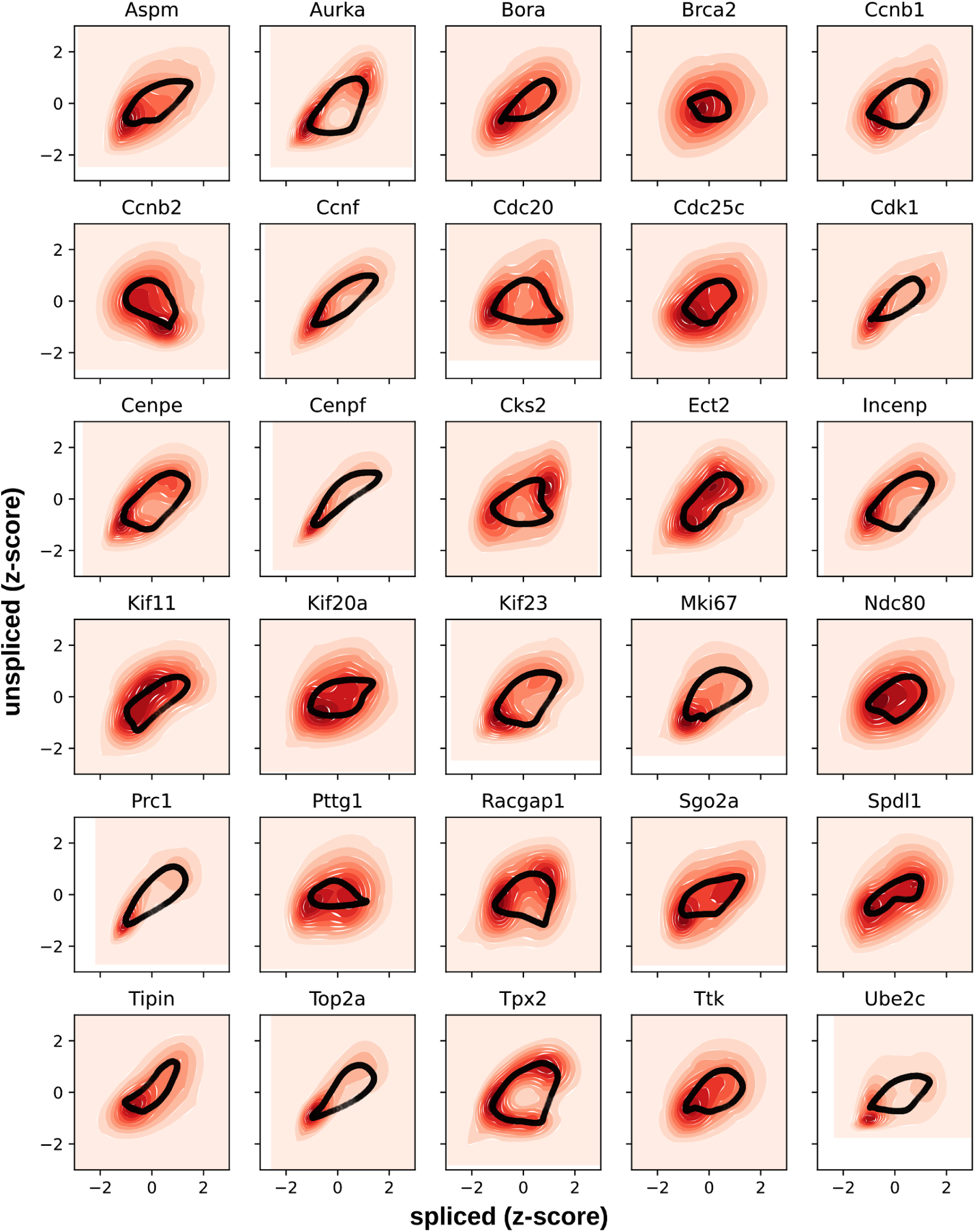
Examples of fits for cycling genes from DeepCycle. Normalized expressions (z-scores) are fed into DeepCycle to extract the circular path.

**Supplementary Figure S5.**
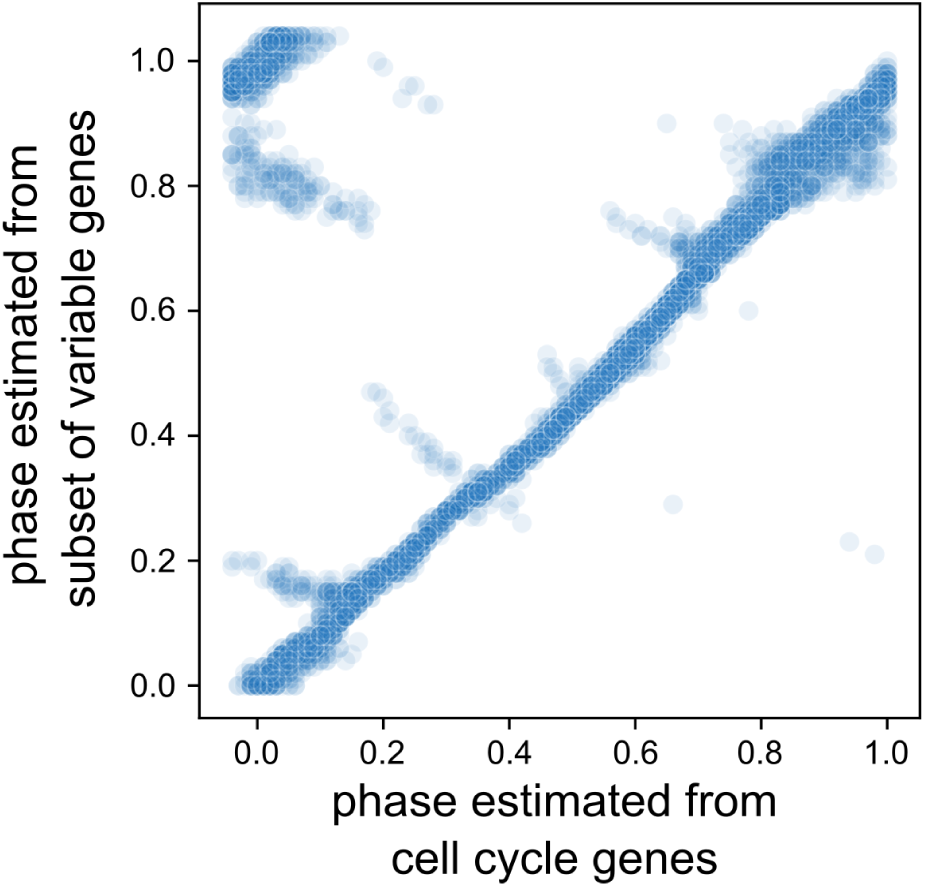
Transcriptional phases are estimated from the selected cell cycle genes (x-axis) and the set of highly variable genes (y-axis). The two phases are highly correlated, showing the robustness of the results even with a lower number of genes.

**Supplementary Figure S6.**
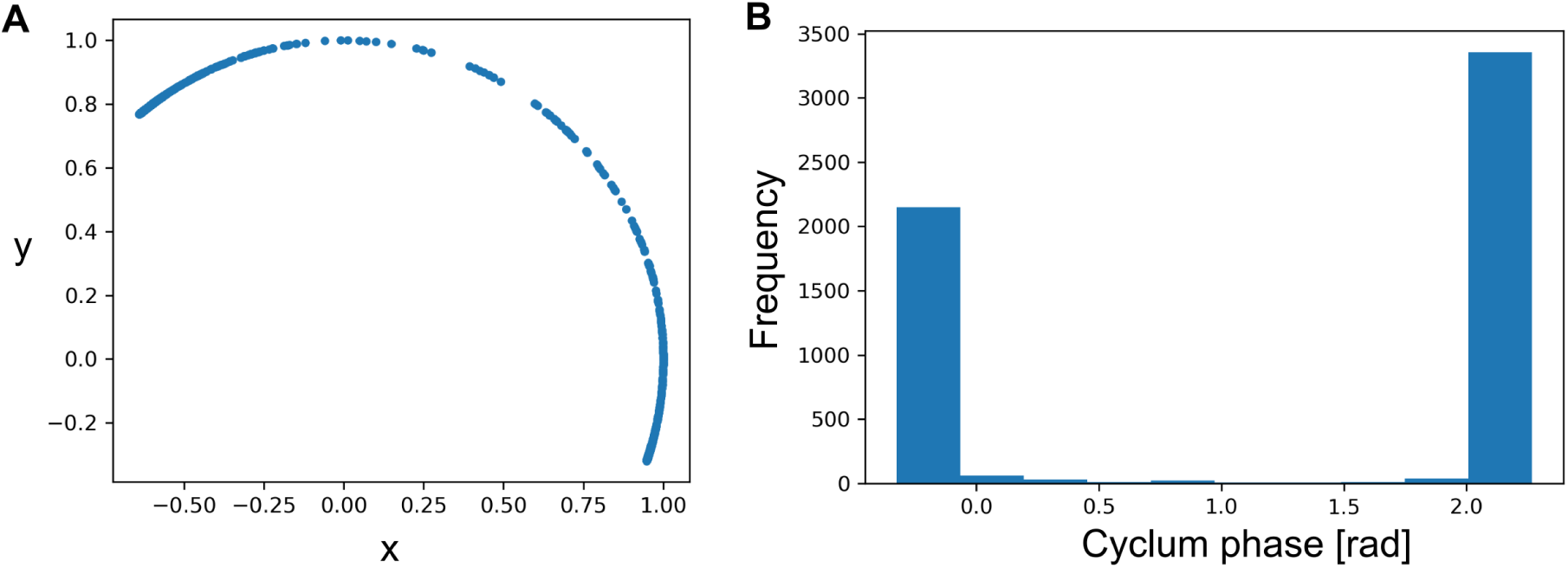
Cyclum^7^ is unable to find the correct cell cycle phase in our data. The angle is not filled by the cells that instead tend to be localized at the opposite sides of the circle.

**Supplementary Figure S7.**
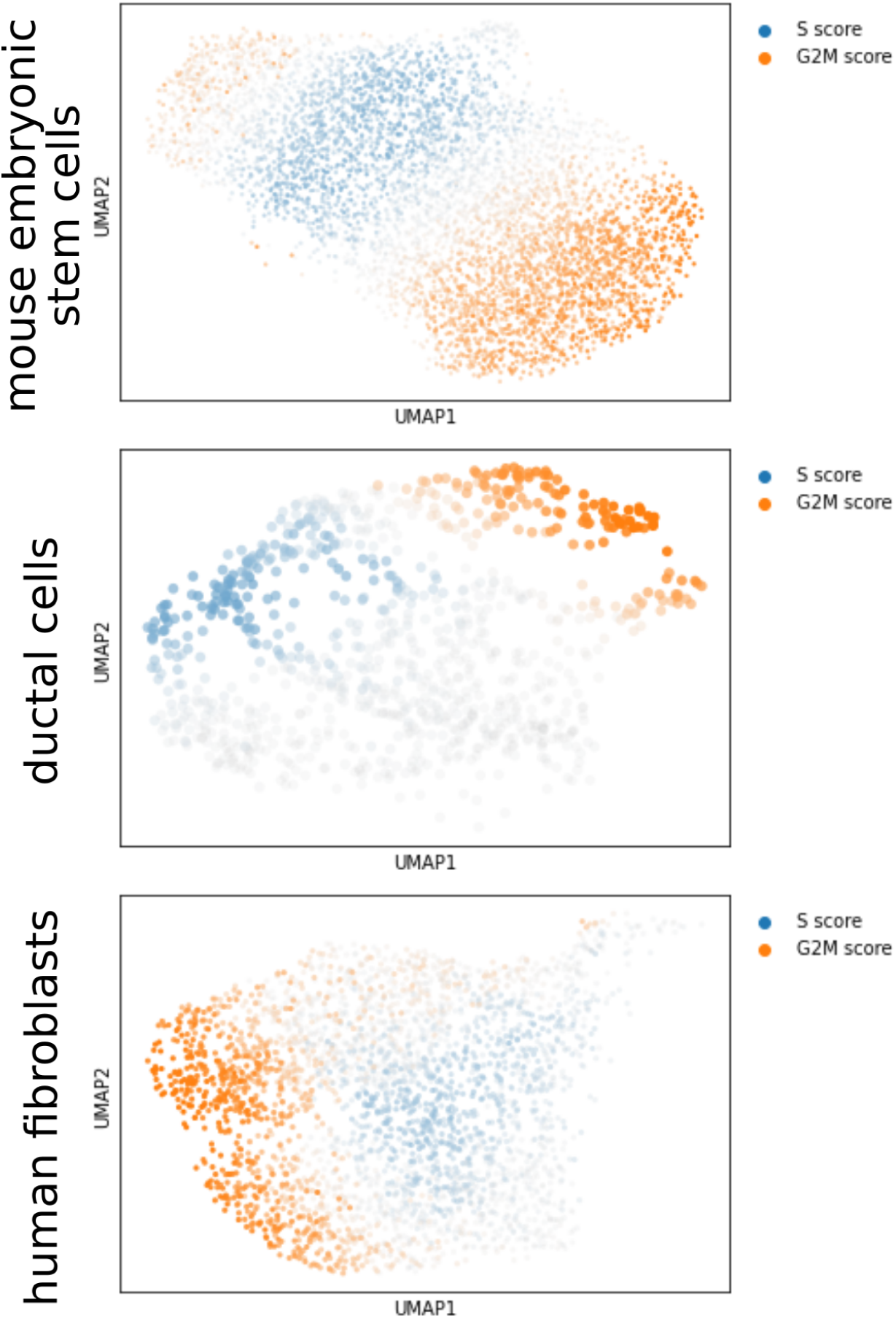
S and G2/M scores for mESCs, ductal cells, and human fibroblasts were calculated from lists of phase-specific genes defined in^19, 29^.

**Supplementary Figure S8.**
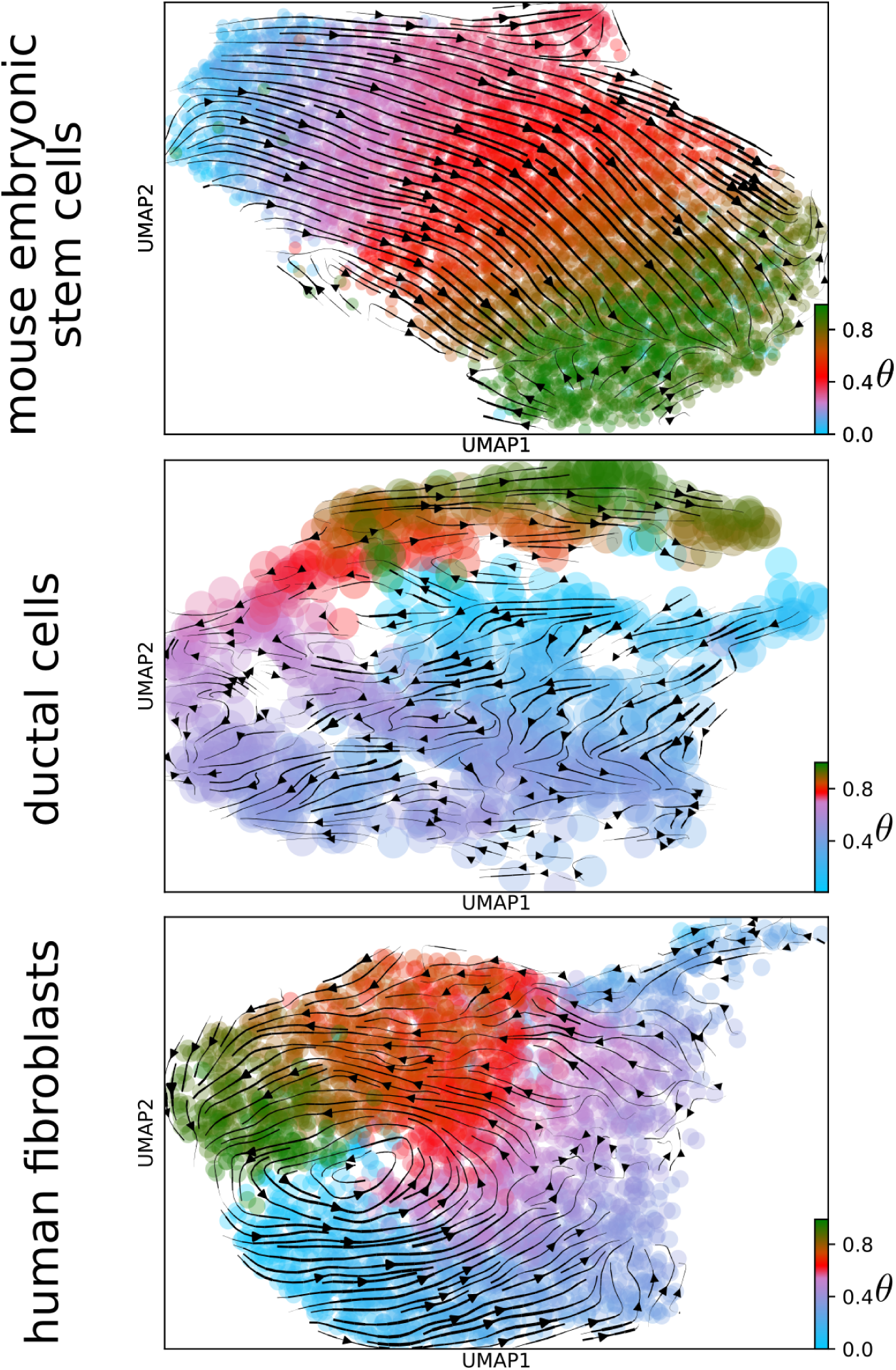
RNA velocity analysis for the mESCs, ductal cells, and the proliferative human fibroblasts.

**Supplementary Figure S9.**
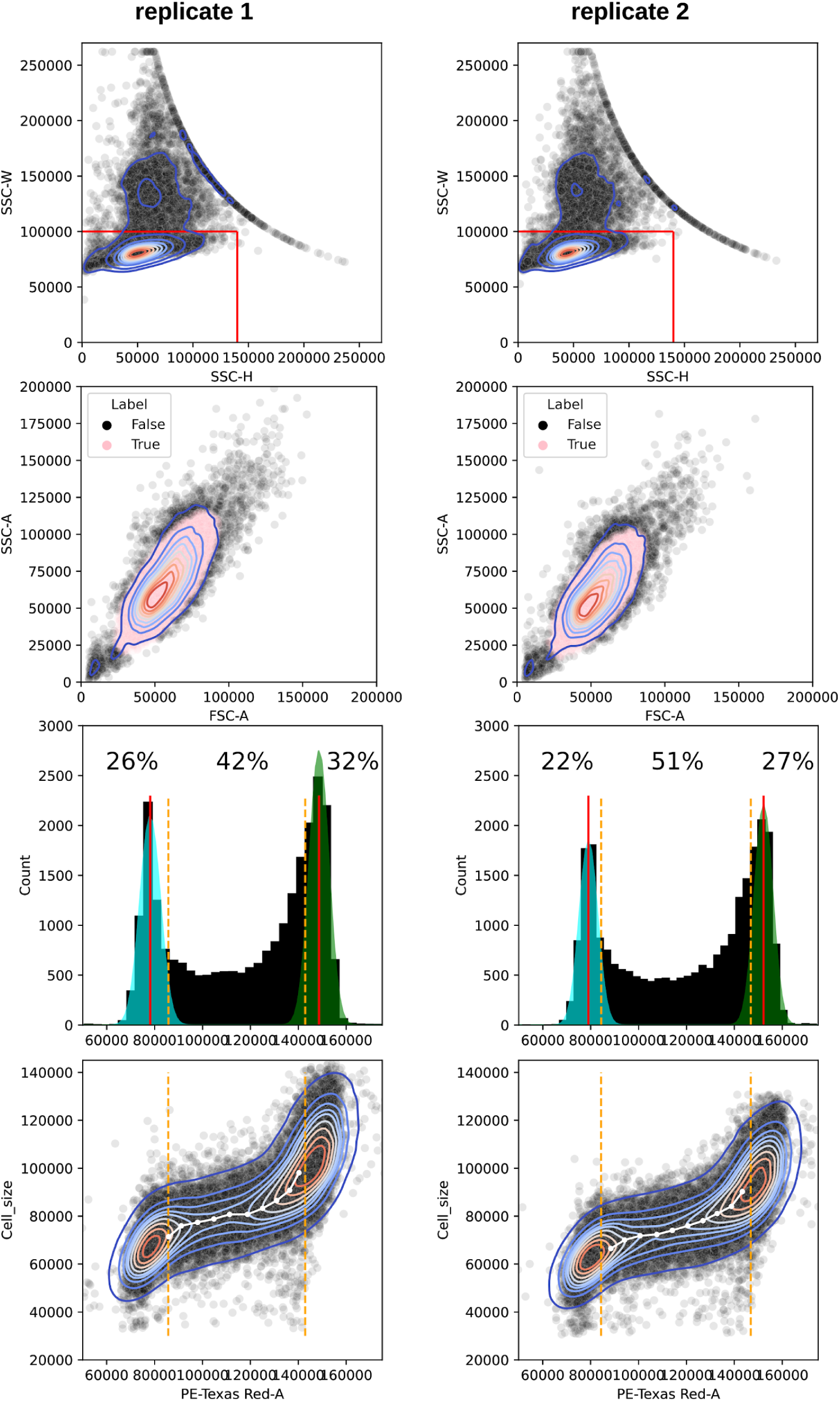
Flow cytometry analysis. The true label shows the cells with the Hotelling T^2^ lower than 6. Orange dashed lines delimit the G1/S and S/G2 transitions.

**Supplementary Figure S10.**
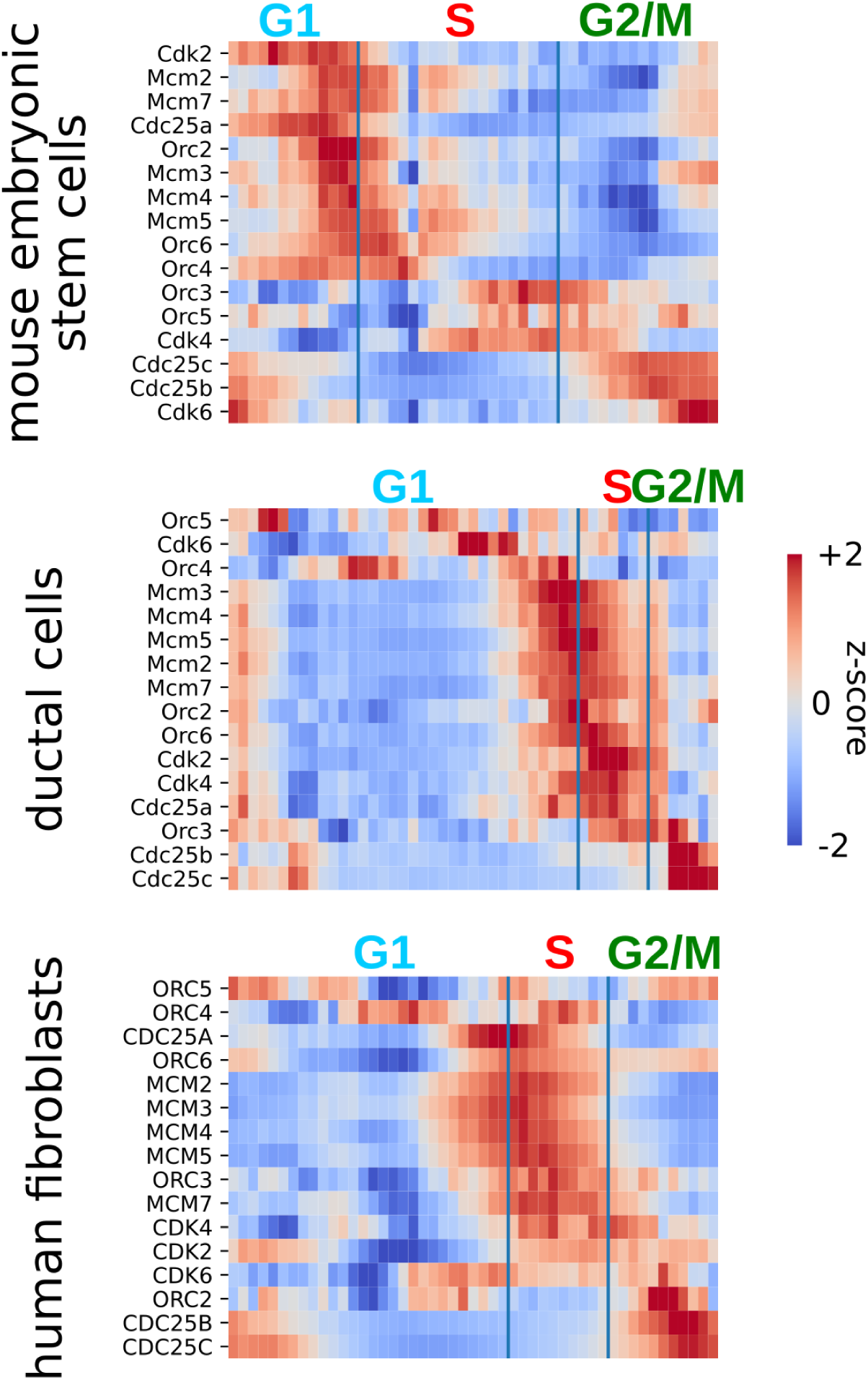
Comparison of core cell cycle genes across datasets.

**Supplementary Figure S11.**
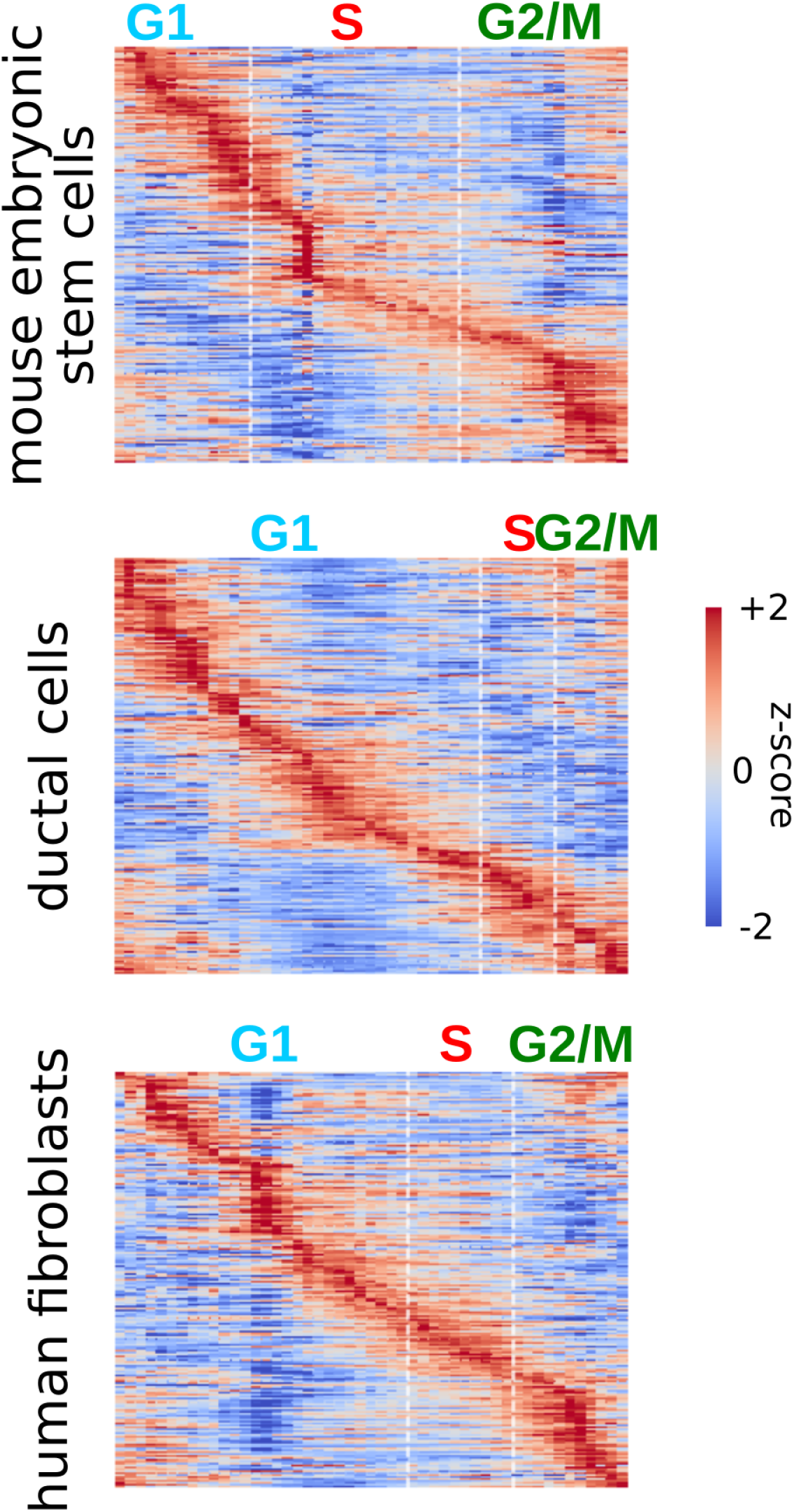
Top genes selected based on their variability in expression across the cell cycle. 4216 genes are shown for mESCs, 1014 for ductal cells, and 3973 for human fibroblasts.

**Supplementary Figure S12.**
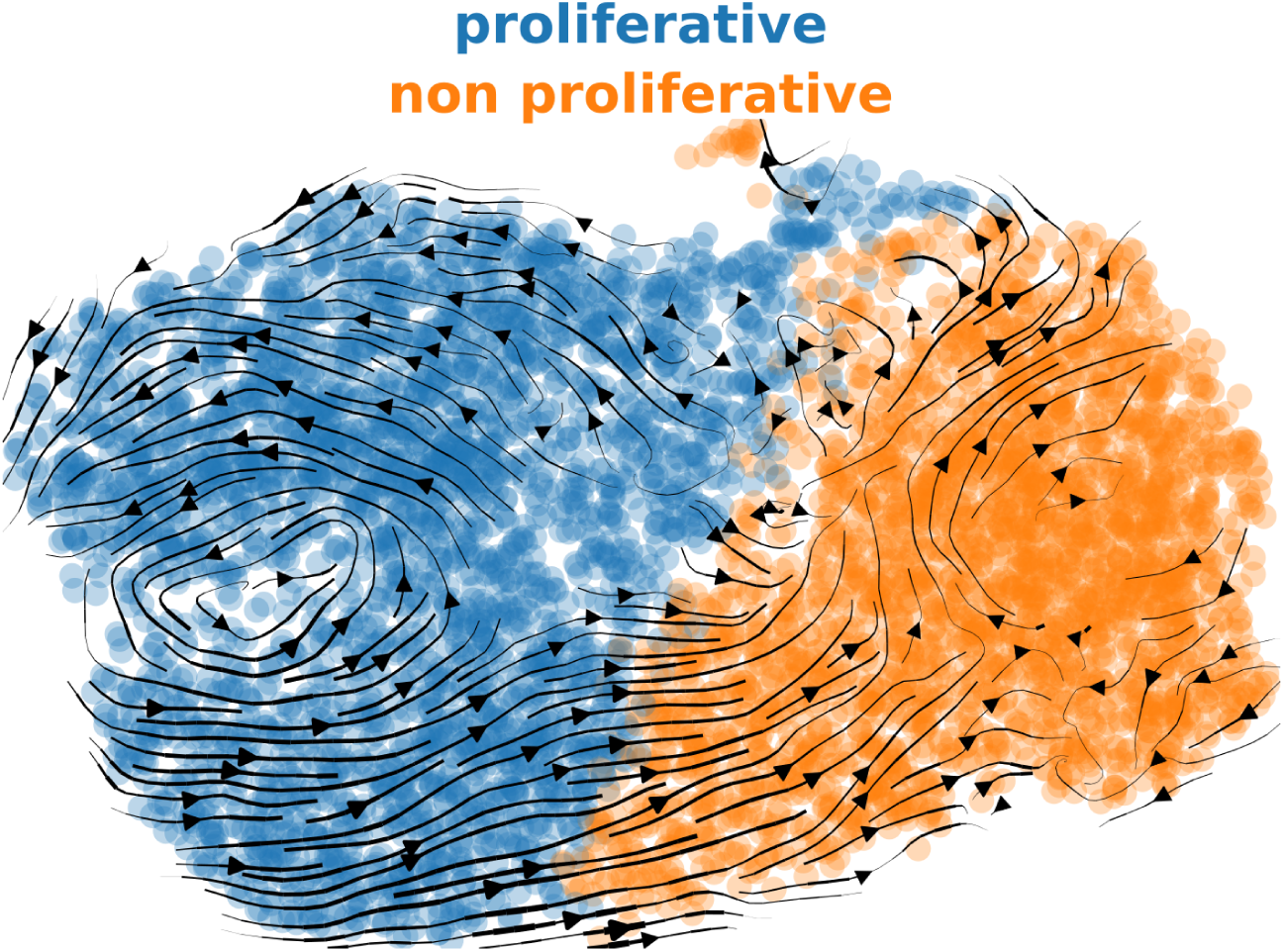
RNA velocity analysis on the whole population of human fibroblasts.

**Supplementary Figure S13.**
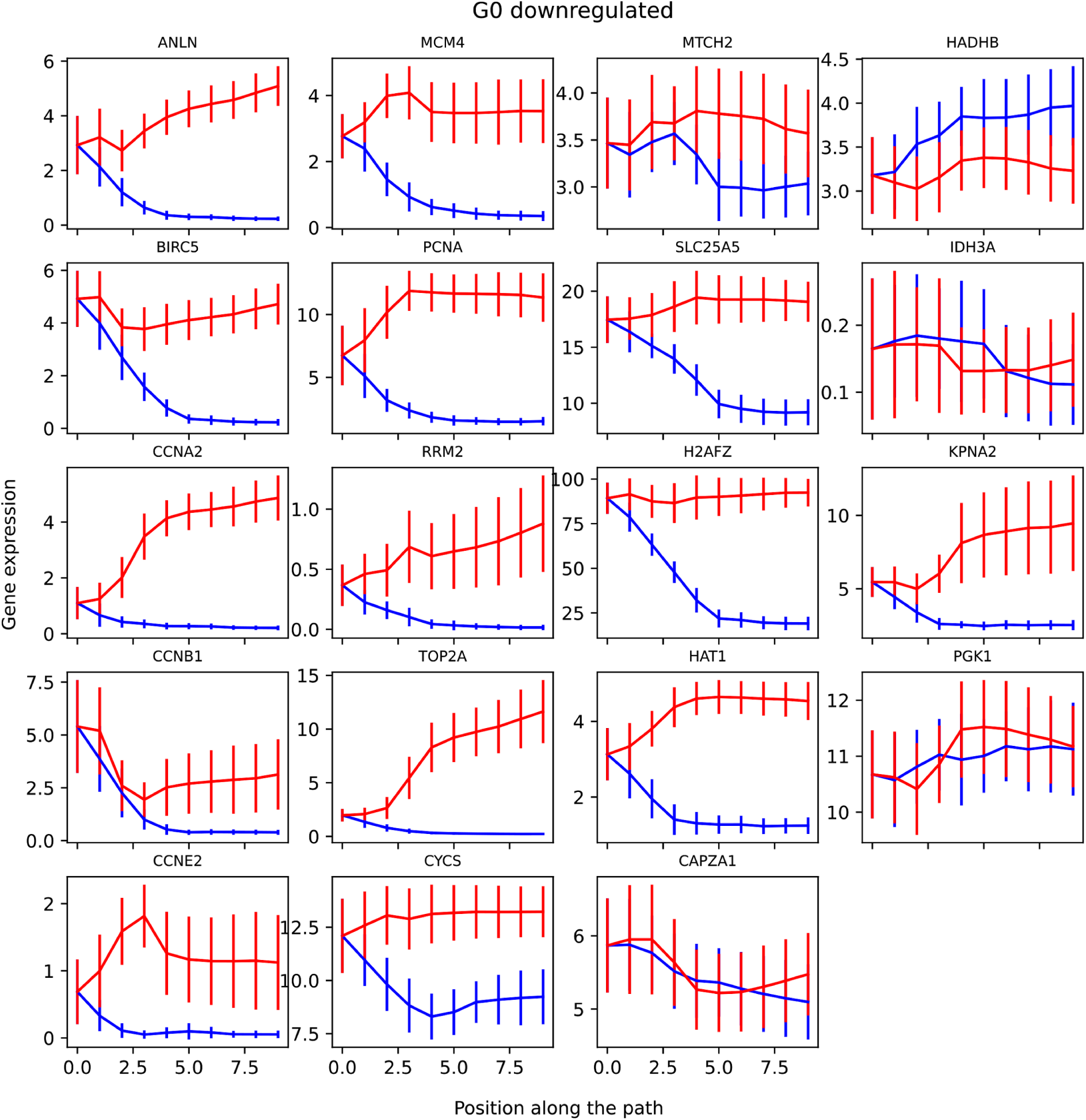
G0 downregulated markers in^62^. The x-axis represents the steps in the paths with the corresponding colors in Figure 5E.

**Supplementary Figure S14.**
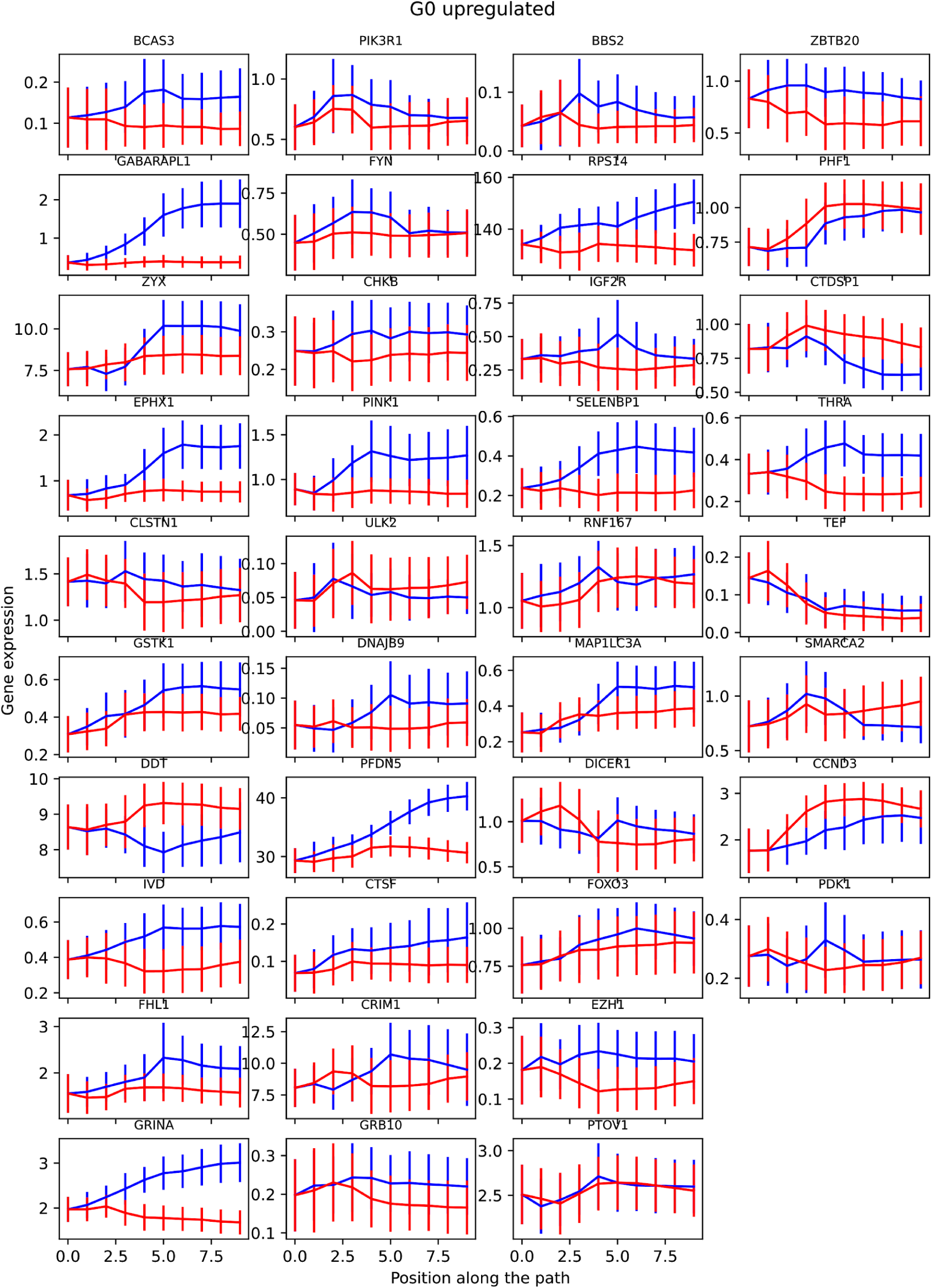
G0 upregulated markers in^62^. The x-axis represents the steps in the paths with the corresponding colors in Figure 5E.

**Supplementary Figure S15.**
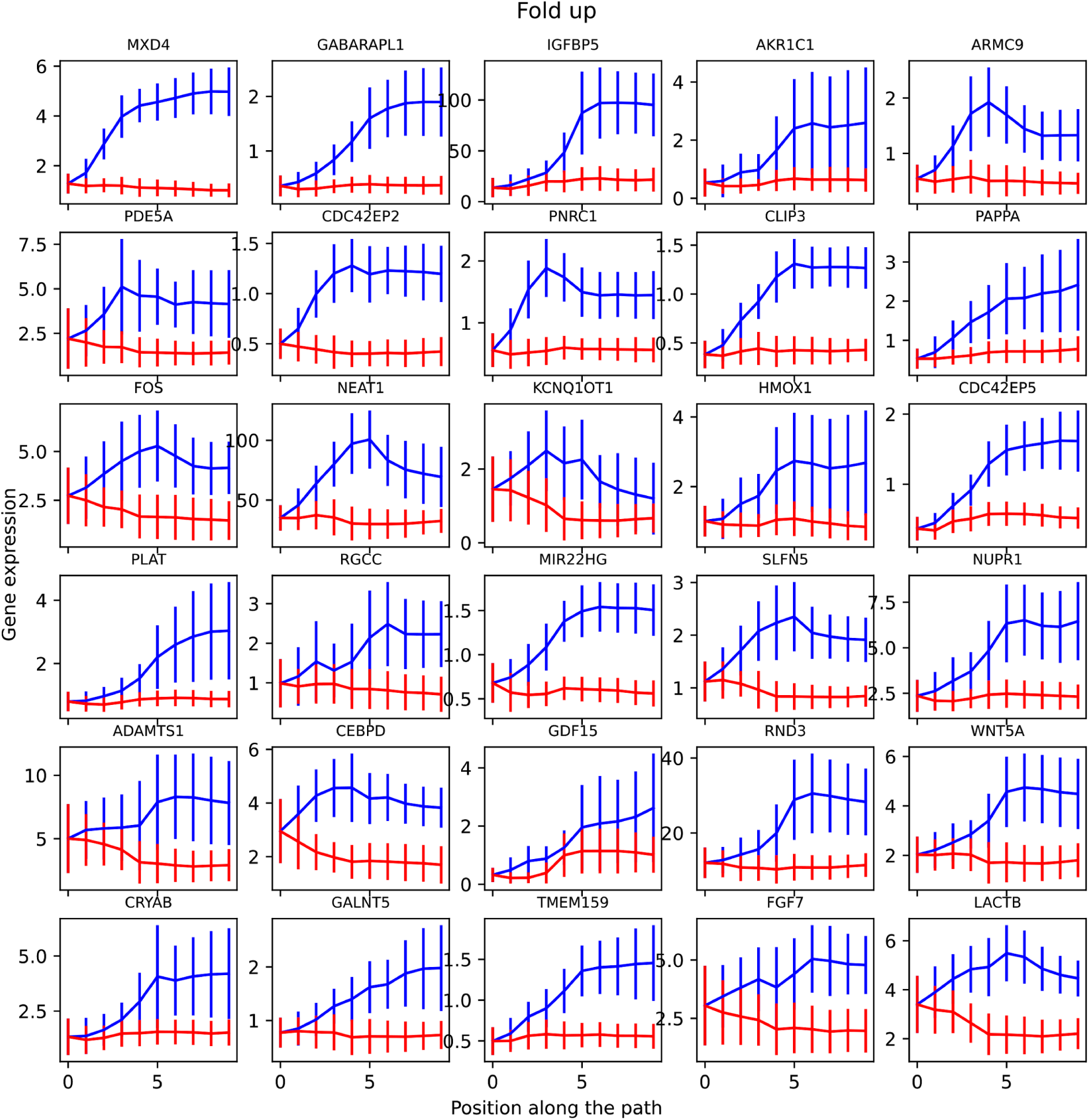
Top genes showing the greatest cumulative fold-up change towards the nonproliferative state. The x-axis represents the steps in the paths with the corresponding colors in Figure 5E.

**Supplementary Figure S16.**
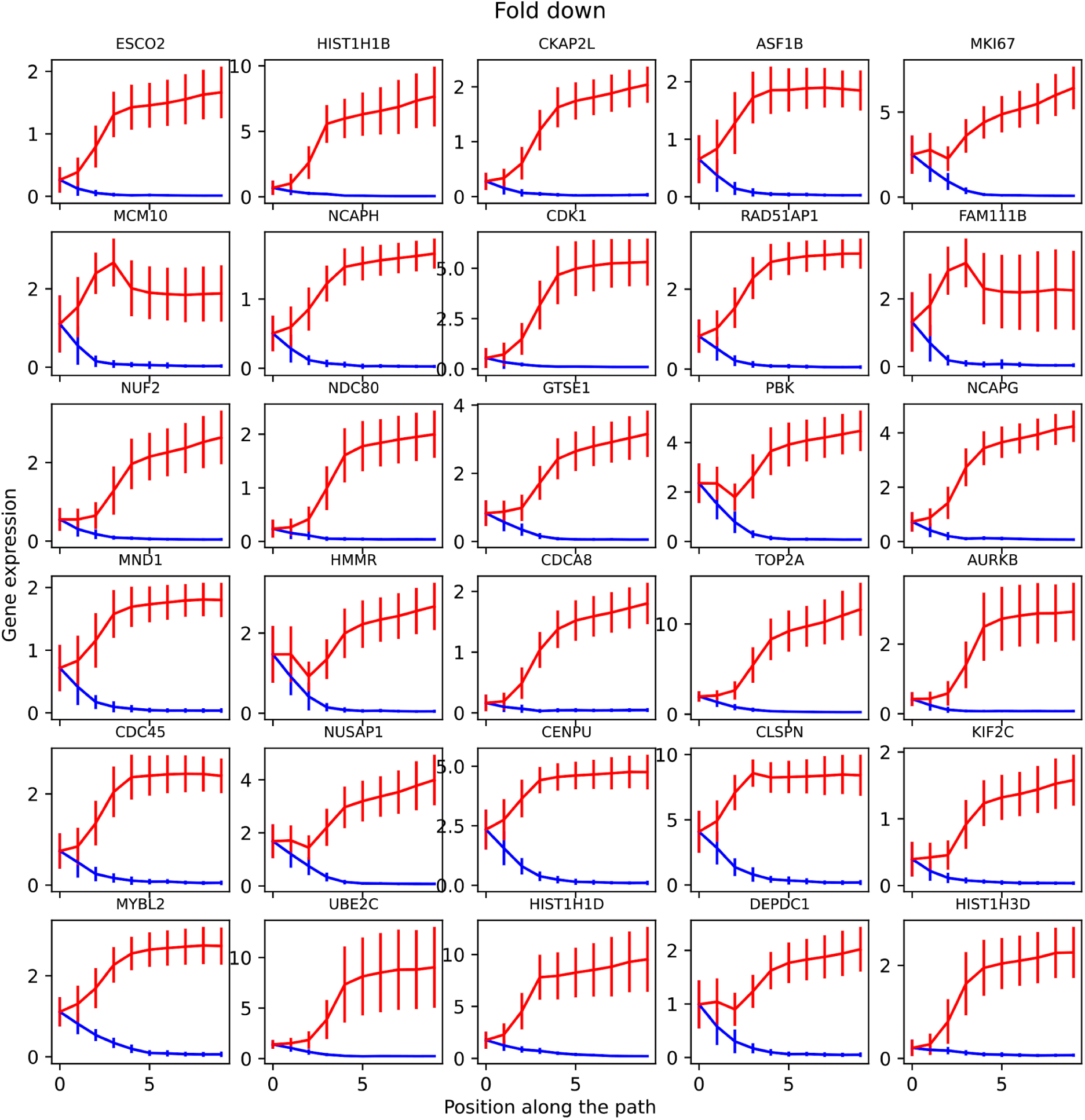
Top genes showing the greatest cumulative fold-down change towards the nonproliferative state. The x-axis represents the steps in the paths with the corresponding colors in Figure 5E.

**Supplementary Figure S17.**
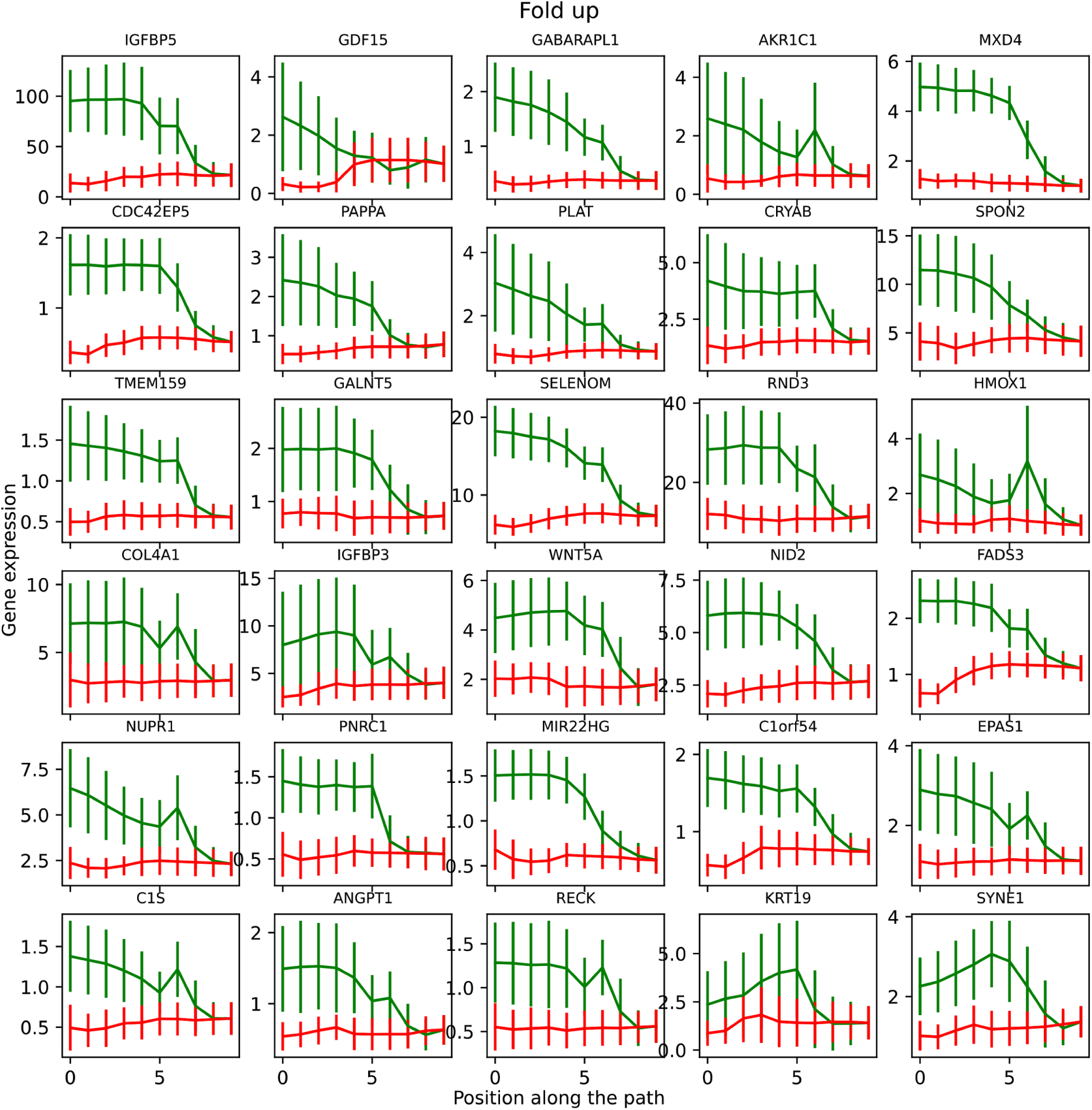
Top genes showing the greatest cumulative fold-up change towards the S phase. The x-axis represents the steps in the paths with the corresponding colors in Figure 5E.

**Supplementary Figure S18.**
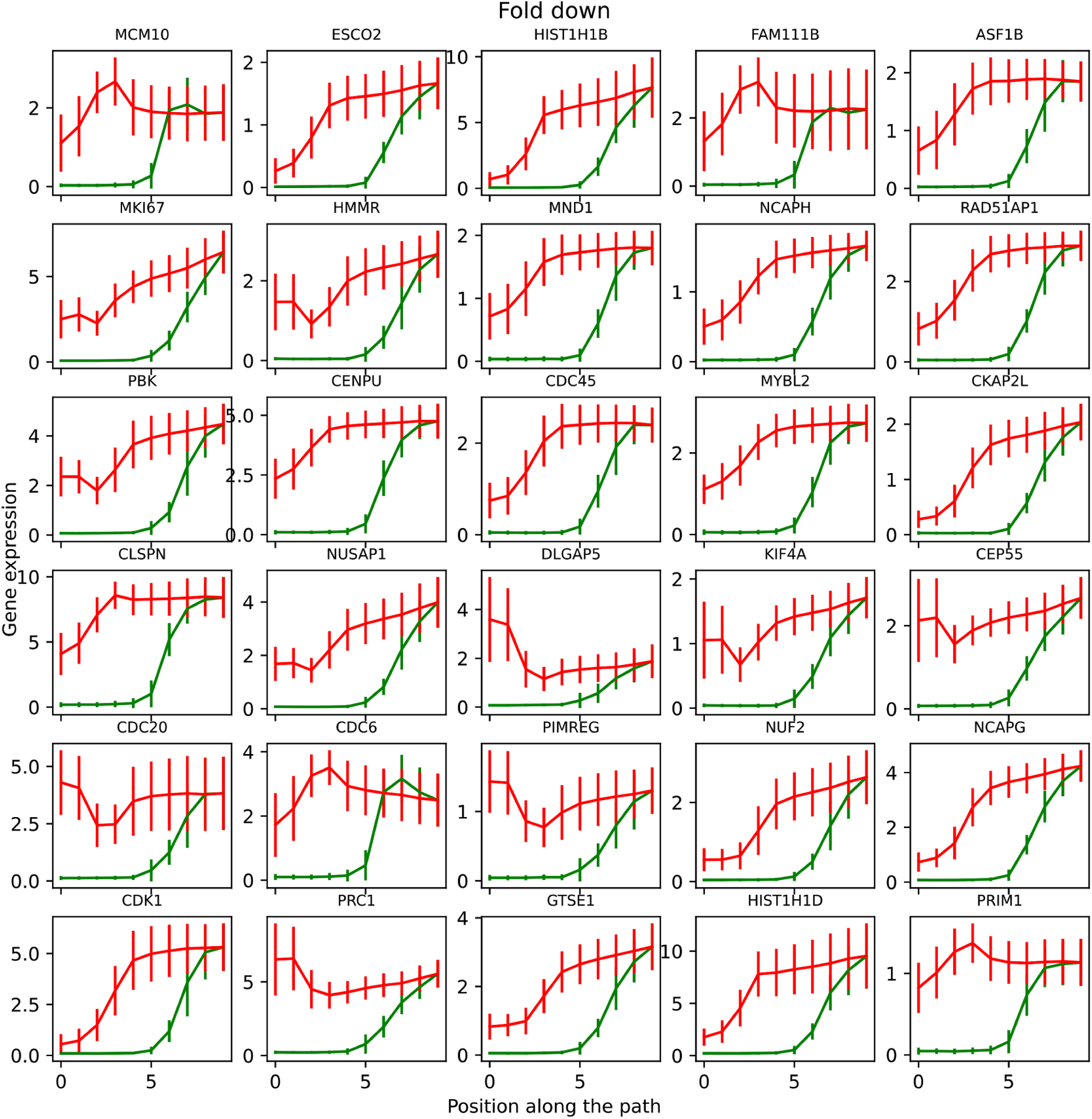
Top genes showing the greatest cumulative fold-down change towards the S phase. The x-axis represents the steps in the paths with the corresponding colors in Figure 5E.

**Supplementary Figure S19.**
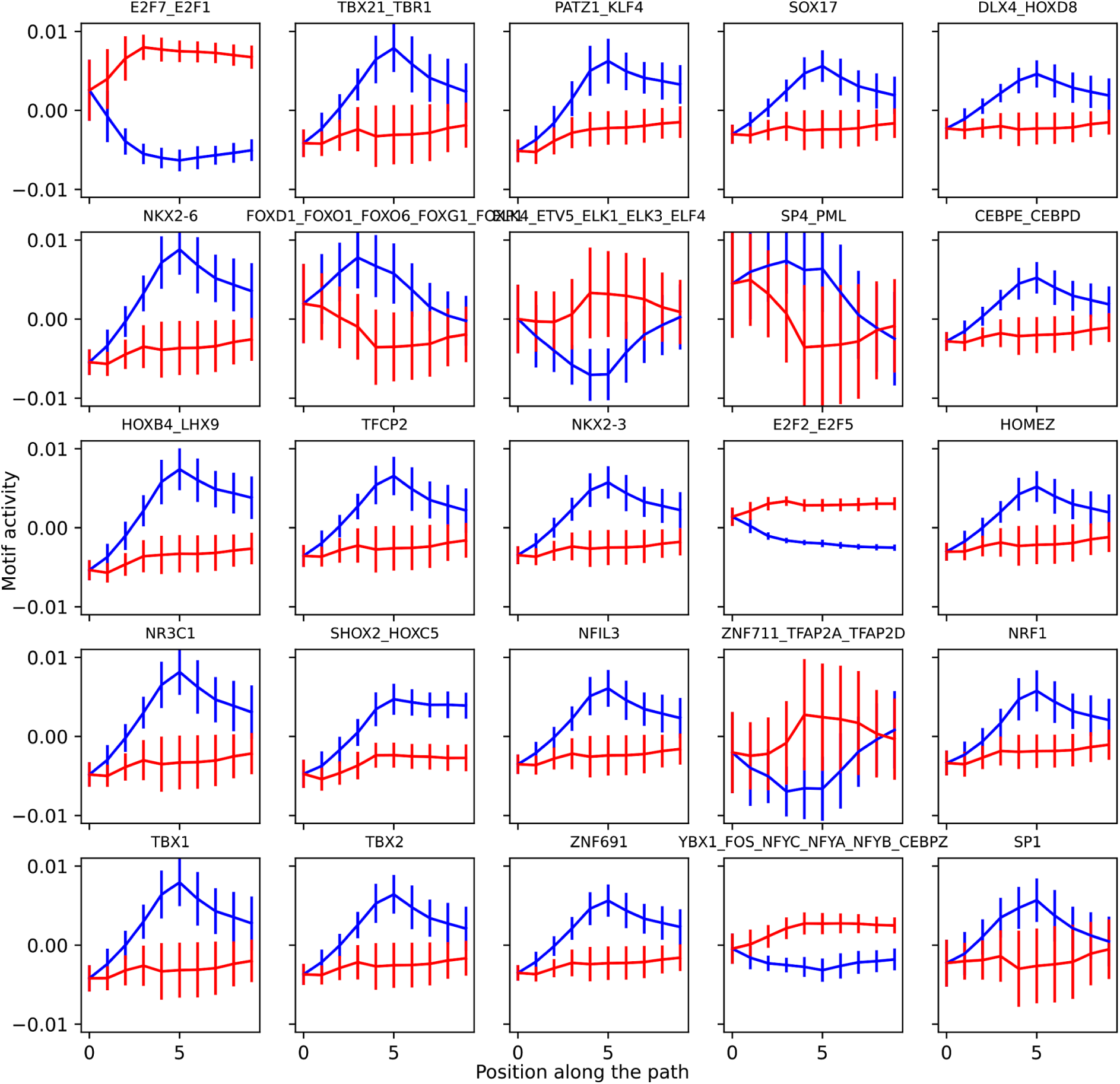
Most significant motifs distinguishing the paths from G1. The x-axis represents the steps in the paths with the corresponding colors in Figure 5E.

**Supplementary Figure S20.**
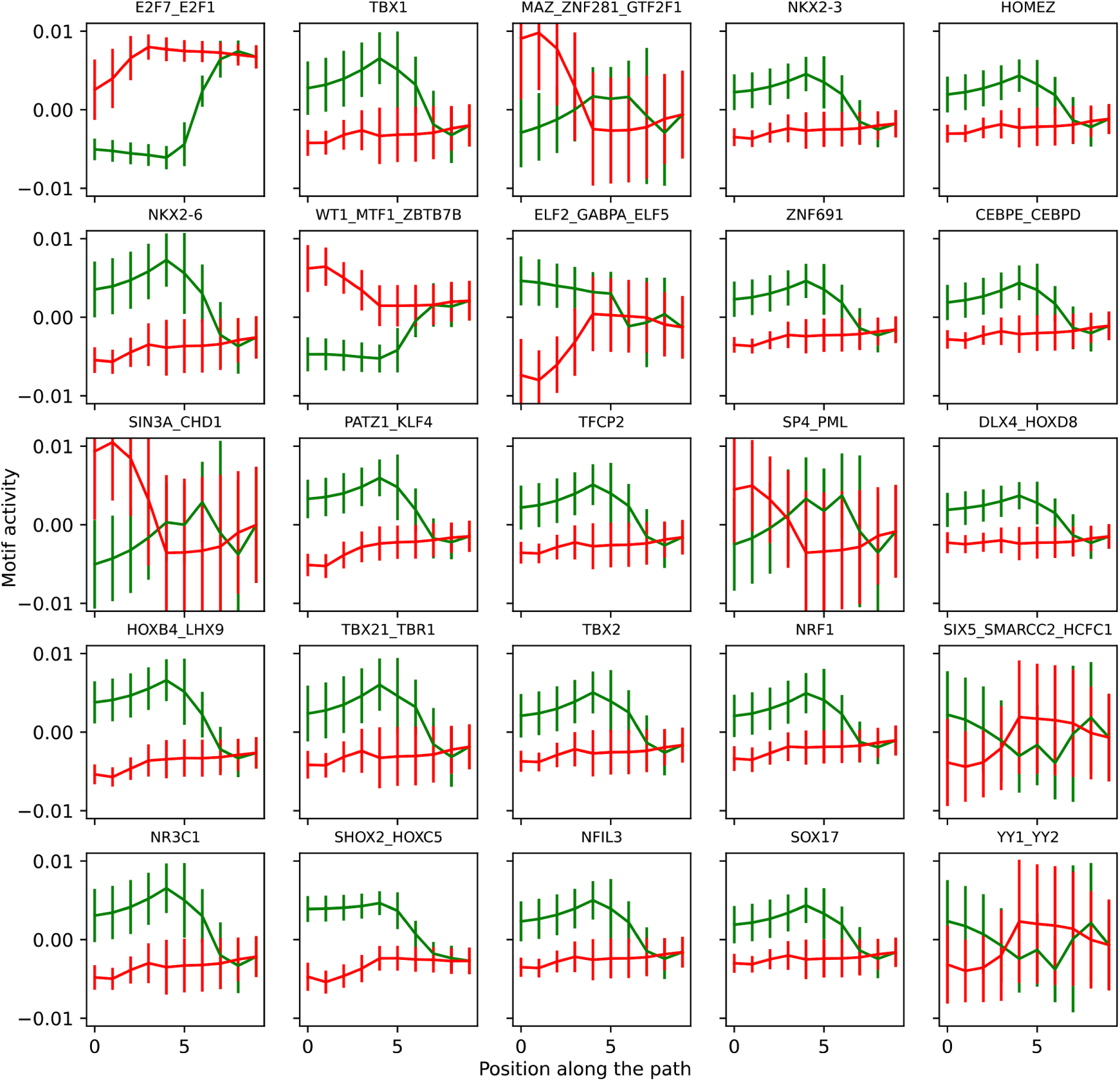
Most significant motifs distinguishing the paths towards S. The x-axis represents the steps in the paths with the corresponding colors in Figure 5E.

## References

1. Alberts, B. et al. Molecular Biology of the Cell. (2007) doi:10.1201/9780203833445.

2. Pardee, A. B. A restriction point for control of normal animal cell proliferation. Proc. Natl. Acad. Sci. U. S. A. 71, 1286–1290 (1974).

3. Hsiao, C. J. et al. Characterizing and inferring quantitative cell cycle phase in single-cell RNA-seq data analysis. Genome Res. 30, 611–621 (2020).

4. Sakaue-Sawano, A. et al. Visualizing spatiotemporal dynamics of multicellular cell-cycle progression. Cell 132, 487–498 (2008).

5. Liu, Z. et al. Reconstructing cell cycle pseudo time-series via single-cell transcriptome data. Nat. Commun. 8, 1–9 (2017).

6. Xia, C., Fan, J., Emanuel, G., Hao, J. & Zhuang, X. Spatial transcriptome profiling by MERFISH reveals subcellular RNA compartmentalization and cell cycle-dependent gene expression. Proc. Natl. Acad. Sci. U. S. A. 116, 19490–19499 (2019).

7. Liang, S., Wang, F., Han, J. & Chen, K. Latent periodic process inference from single-cell RNA-seq data. Nat. Commun. 11, 1–8 (2020).

8. Schwabe, D., Formichetti, S., Junker, J. P., Falcke, M. & Rajewsky, N. The transcriptome dynamics of single cells during the cell cycle. Mol. Syst. Biol. 16, e9946 (2020).

9. La Manno, G. et al. RNA velocity of single cells. Nature 560, 494–498 (2018).

10. Pauklin, S. & Vallier, L. The Cell-Cycle State of Stem Cells Determines Cell Fate Propensity. Cell 156, 1338 (2014).

11. Dalton, S. Linking the Cell Cycle to Cell Fate Decisions. Trends Cell Biol. 25, 592–600 (2015).

12. Jirawatnotai, S., Dalton, S. & Wattanapanitch, M. Role of cyclins and cyclin-dependent kinases in pluripotent stem cells and their potential as a therapeutic target. Semin. Cell Dev. Biol. 107, 63–71 (2020).

13. Ruiz, S. et al. A high proliferation rate is required for cell reprogramming and maintenance of human embryonic stem cell identity. Curr. Biol. 21, 45–52 (2011).

14. Matson, J. P. & Cook, J. G. Cell cycle proliferation decisions: the impact of single cell analyses. FEBS J. 284, 362–375 (2017).

15. Coronado, D. et al. A short G1 phase is an intrinsic determinant of naïve embryonic stem cell pluripotency. Stem Cell Research vol. 10 118–131 (2013).

16. Nichols, J. & Smith, A. Naive and Primed Pluripotent States. Cell Stem Cell vol. 4 487–492 (2009).

17. Zaveri, L. & Dhawan, J. Cycling to Meet Fate: Connecting Pluripotency to the Cell Cycle. Frontiers in Cell and Developmental Biology vol. 6 (2018).

18. Bastidas-Ponce, A. et al. Comprehensive single cell mRNA profiling reveals a detailed roadmap for pancreatic endocrinogenesis. Development 146, (2019).

19. Bergen, V., Lange, M., Peidli, S., Wolf, F. A. & Theis, F. J. Generalizing RNA velocity to transient cell states through dynamical modeling. Nat. Biotechnol. (2020) doi:10.1038/s41587-020-0591-3.

20. Kratsios, A. The Universal Approximation Property. Annals of Mathematics and Artificial Intelligence (2021) doi:10.1007/s10472-020-09723-1.

21. Lopez, R., Regier, J., Cole, M. B., Jordan, M. I. & Yosef, N. Deep generative modeling for single-cell transcriptomics. Nat. Methods 15, 1053–1058 (2018).

22. Grønbech, C. H. et al. scVAE: Variational auto-encoders for single-cell gene expression data. Bioinformatics (2020) doi:10.1093/bioinformatics/btaa293.

23. Eraslan, G., Simon, L. M., Mircea, M., Mueller, N. S. & Theis, F. J. Single-cell RNA-seq denoising using a deep count autoencoder. Nat. Commun. 10, 390 (2019).

24. Talwar, D., Mongia, A., Sengupta, D. & Majumdar, A. AutoImpute: Autoencoder based imputation of single-cell RNA-seq data. Sci. Rep. 8, 16329 (2018).

25. Wang, D. & Gu, J. VASC: Dimension Reduction and Visualization of Single-cell RNA-seq Data by Deep Variational Autoencoder. Genomics, Proteomics & Bioinformatics vol. 16 320–331 (2018).

26. Deng, Y., Bao, F., Dai, Q., Wu, L. F. & Altschuler, S. J. Scalable analysis of cell-type composition from single-cell transcriptomics using deep recurrent learning. Nat. Methods 16, 311–314 (2019).

27. Ding, J., Condon, A. & Shah, S. P. Interpretable dimensionality reduction of single cell transcriptome data with deep generative models. Nat. Commun. 9, 2002 (2018).

28. Wang, J. et al. Data denoising with transfer learning in single-cell transcriptomics. Nat. Methods 16, 875–878 (2019).

29. Tirosh, I. et al. Dissecting the multicellular ecosystem of metastatic melanoma by single-cell RNA-seq. Science 352, 189–196 (2016).

30. Kellogg, D. R. Wee1-dependent mechanisms required for coordination of cell growth and cell division. J. Cell Sci. 116, 4883–4890 (2003).

31. Kim, S. Y. & Ferrell, J. E., Jr. Substrate competition as a source of ultrasensitivity in the inactivation of Wee1. Cell 128, 1133–1145 (2007).

32. Marumoto, T. et al. Aurora-A kinase maintains the fidelity of early and late mitotic events in HeLa cells. J. Biol. Chem. 278, 51786–51795 (2003).

33. Honda, R., Körner, R. & Nigg, E. A. Exploring the functional interactions between Aurora B, INCENP, and survivin in mitosis. Mol. Biol. Cell 14, 3325–3341 (2003).

34. Chou, H.-Y. et al. Phosphorylation of NuSAP by Cdk1 regulates its interaction with microtubules in mitosis. Cell Cycle 10, 4083–4089 (2011).

35. Li, C. et al. NuSAP modulates the dynamics of kinetochore microtubules by attenuating MCAK depolymerisation activity. Sci. Rep. 6, 18773 (2016).

36. Liu, K. et al. The role of CDC25C in cell cycle regulation and clinical cancer therapy: a systematic review. Cancer Cell Int. 20, 1–16 (2020).

37. Sur, S. & Agrawal, D. K. Phosphatases and kinases regulating CDC25 activity in the cell cycle: clinical implications of CDC25 overexpression and potential treatment strategies. Mol. Cell. Biochem. 416, 33–46 (2016).

38. Shen, T. & Huang, S. The Role of Cdc25A in the Regulation of Cell Proliferation and Apoptosis. Anticancer Agents Med. Chem. 12, 631–639 (2012).

39. Hoffmann, I. The role of Cdc25 phosphatases in cell cycle checkpoints. Protoplasma 211, 8–11 (2000).

40. Bochman, M. L. & Schwacha, A. The Mcm complex: unwinding the mechanism of a replicative helicase. Microbiol. Mol. Biol. Rev. 73, 652–683 (2009).

41. Meijer, L. et al. Biochemical and cellular effects of roscovitine, a potent and selective inhibitor of the cyclin-dependent kinases cdc2, cdk2 and cdk5. Eur. J. Biochem. 243, 527–536 (1997).

42. Liu, L., Michowski, W., Kolodziejczyk, A. & Sicinski, P. The cell cycle in stem cell proliferation, pluripotency and differentiation. Nat. Cell Biol. 21, 1060–1067 (2019).

43. Balwierz, P. J. et al. ISMARA: automated modeling of genomic signals as a democracy of regulatory motifs. Genome Res. 24, 869–884 (2014).

44. Wang, J. et al. YY1 Positively Regulates Transcription by Targeting Promoters and Super-Enhancers through the BAF Complex in Embryonic Stem Cells. Stem Cell Reports vol. 10 1324–1339 (2018).

45. Raccaud, M. et al. Mitotic chromosome binding predicts transcription factor properties in interphase. Nat. Commun. 10, 487 (2019).

46. Tsai, S.-Y. et al. Mouse development with a single E2F activator. Nature 454, 1137–1141 (2008).

47. Gaubatz, S. et al. E2F4 and E2F5 Play an Essential Role in Pocket Protein–Mediated G1 Control. Molecular Cell vol. 6 729–735 (2000).

48. Timmers, C. et al. E2f1, E2f2, and E2f3 Control E2F Target Expression and Cellular Proliferation via a p53-Dependent Negative Feedback Loop. Mech. Chem. Biosyst. 27, 65–78 (2007).

49. Kotake, Y., Arikawa, N., Tahara, K., Maru, H. & Naemura, M. Y-box binding protein 1 is involved in regulating the g2/m phase of the cell cycle. Anticancer Res. 37, 1603–1608 (2017).

50. Jurchott, K. et al. YB-1 as a cell cycle-regulated transcription factor facilitating cyclin A and cyclin B1 gene expression. J. Biol. Chem. 278, 27988–27996 (2003).

51. Liu, Z. et al. Overexpression of YBX1 Promotes Pancreatic Ductal Adenocarcinoma Growth via the GSK3B/Cyclin D1/Cyclin E1 Pathway. Mol Ther Oncolytics 17, 21–30 (2020).

52. Nakata, Y. et al. c-Myb contributes to G2/M cell cycle transition in human hematopoietic cells by direct regulation of cyclin B1 expression. Mol. Cell. Biol. 27, 2048–2058 (2007).

53. Nakata, Y. et al. c-Myb Plays a Role InG2/M Cell Cycle Transition by Direct Regulation of Cyclin B1 Expression in Hematopoietic Cells. Blood vol. 106 1355–1355 (2005).

54. Álvaro-Blanco, J. et al. MAZ induces MYB expression during the exit from quiescence via the E2F site in the MYB promoter. Nucleic Acids Res. 45, 9960–9975 (2017).

55. Laoukili, J., Stahl, M. & Medema, R. H. FoxM1: at the crossroads of ageing and cancer. Biochim. Biophys. Acta 1775, 92–102 (2007).

56. Liao, G.-B. et al. Regulation of the master regulator FOXM1 in cancer. Cell Commun. Signal. 16, 57 (2018).

57. Dunn, S.-. J., Martello, G., Yordanov, B., Emmott, S. & Smith, A. G. Defining an essential transcription factor program for naive pluripotency. Science vol. 344 1156–1160 (2014).

58. Matsuda, T. et al. STAT3 activation is sufficient to maintain an undifferentiated state of mouse embryonic stem cells. EMBO J. 18, 4261–4269 (1999).

59. Ghaleba, A. M. & Yang, V. W. Krüppel-like factor 4 (KLF4): What we currently know. Gene 611, 27–37 (2017).

60. Festuccia, N. et al. Transcription factor activity and nucleosome organization in mitosis. Genome Res. 29, 250–260 (2019).

61. Festuccia, N. et al. Mitotic binding of Esrrb marks key regulatory regions of the pluripotency network. Nat. Cell Biol. 18, 1139–1148 (2016).

62. Cheung, T. H. & Rando, T. A. Molecular regulation of stem cell quiescence. Nat. Rev. Mol. Cell Biol. 14, 329–340 (2013).

63. Henkelman, G., Uberuaga, B. P. & Jónsson, H. A climbing image nudged elastic band method for finding saddle points and minimum energy paths. J. Chem. Phys. 113, 9901–9904 (2000).

64. Rognoni, E. et al. Fibroblast state switching orchestrates dermal maturation and wound healing. Mol. Syst. Biol. 14, e8174 (2018).

65. Boros, K., Lacaud, G. & Kouskoff, V. The transcription factor Mxd4 controls the proliferation of the first blood precursors at the onset of hematopoietic development in vitro. Exp. Hematol. 39, 1090–1100 (2011).

66. Farrugia, A. J. & Calvo, F. The Borg family of Cdc42 effector proteins Cdc42EP1–5. Biochem. Soc. Trans. 44, 1709–1716 (2016).

67. Ding, J. & Du, K. ClipR-59 interacts with Akt and regulates Akt cellular compartmentalization. Mol. Cell. Biol. 29, 1459–1471 (2009).

68. Su, W. et al. Silencing of Long Noncoding RNA MIR22HG Triggers Cell Survival/Death Signaling via Oncogenes YBX1, MET, and p21 in Lung Cancer. Cancer Res. 78, 3207–3219 (2018).

69. Han, M. et al. Interfering with long non-coding RNA MIR22HG processing inhibits glioblastoma progression through suppression of Wnt/β-catenin signalling. Brain 143, 512–530 (2020).

70. Alomer, R. M. et al. Esco1 and Esco2 regulate distinct cohesin functions during cell cycle progression. Proc. Natl. Acad. Sci. U. S. A. 114, 9906–9911 (2017).

71. Izumi, M., Yatagai, F. & Hanaoka, F. Cell Cycle-dependent Proteolysis and Phosphorylation of Human Mcm10*. J. Biol. Chem. 276, 48526–48531 (2001).

72. Musa, J., Aynaud, M.-M., Mirabeau, O., Delattre, O. & Grünewald, T. G. MYBL2 (B-Myb): a central regulator of cell proliferation, cell survival and differentiation involved in tumorigenesis. Cell Death Dis. 8, e2895 (2017).

73. Zhang, T. et al. Nuf2 is required for chromosome segregation during mouse oocyte meiotic maturation. Cell Cycle 14, 2701–2710 (2015).

74. Hori, T., Haraguchi, T., Hiraoka, Y., Kimura, H. & Fukagawa, T. Dynamic behavior of Nuf2-Hec1 complex that localizes to the centrosome and centromere and is essential for mitotic progression in vertebrate cells. J. Cell Sci. 116, 3347–3362 (2003).

75. Takahashi, K. & Yamanaka, S. Induction of Pluripotent Stem Cells from Mouse Embryonic and Adult Fibroblast Cultures by Defined Factors. Cell 126, 663–676 (2006).

76. Wolf, F. A., Angerer, P. & Theis, F. J. SCANPY: large-scale single-cell gene expression data analysis. Genome Biol. 19, 15 (2018).

77. Arnold, P., Erb, I., Pachkov, M., Molina, N. & van Nimwegen, E. MotEvo: integrated Bayesian probabilistic methods for inferring regulatory sites and motifs on multiple alignments of DNA sequences. Bioinformatics 28, 487–494 (2012).

